# Flexibility and modulation of translation initiation in enterovirus genomes

**DOI:** 10.1101/2025.03.24.645098

**Authors:** Rhian L. O’Connor, Georgia M. Cook, Jacqueline Hankinson, Ksenia Fominykh, Samantha H. Cheng, Daniel A. Nash, Aurélie Cenier, Komal M. Nayak, Stephen C. Graham, Janet E. Deane, Matthias Zilbauer, Andrew E. Firth, Valeria Lulla

**Author notes:** Correspondence to (VL).

## Abstract

Enteroviruses comprise a large group of mammalian pathogens that often utilize two open reading frames (ORFs) to encode their proteins: the upstream protein (UP) and the main polyprotein. In some enteroviruses, in addition to the canonical upstream AUG (uAUG), there is another AUG that may represent an alternative upstream initiation site. An analysis of enterovirus sequences containing additional upstream AUGs identified several clusters, including strains of pathogenic *Enterovirus alphacoxsackie* and *E. coxsackiepol*. Using ribosome profiling on coxsackievirus CVA-13 (*E. coxsackiepol*), we demonstrate that both upstream AUG codons can be used for translation initiation in infected cells. Moreover, we confirm translation from both upstream AUGs using a reporter system. Mutating the additional upstream AUG in the context of CVA-13 did not result in phenotypic changes in immortalized cell lines. However, the wild-type virus outcompeted this mutant in human intestinal organoids and differentiated neuronal systems, representing an advantage in physiologically relevant infection sites. Mutation of the stop codon of the shorter upstream ORF led to dysregulated translation of the other ORFs in the reporter system, suggesting a potential role for the additional uORF in modulating the expression level of the other ORFs. These findings demonstrate the remarkable plasticity of enterovirus IRES-mediated initiation and the competitive advantage of double-upstream-AUG-containing viruses in terminally differentiated intestinal organoids and neuronal systems.

## Introduction

The *Enterovirus* genus belongs to the *Picornaviridae* family and consists of 13 species and more than 70 serotypes. Human enteroviruses are widely spread respiratory and intestinal pathogens and are classified into four species: *Enterovirus alphacoxsackie, E. betacoxsackie, E. coxsackiepol* and *E. deconjuncti* (formerly *Enterovirus A* to *D*). Virus infections usually begin in peripheral tissues and can progress to the nervous system. Disease phenotype in humans ranges from sub-clinical to acute flaccid paralysis, myocarditis, and meningitis ^1–4^. Enteroviruses are non-enveloped viruses with single-stranded positive-sense RNA genomes of ∼7.4 kb (Fig. 1A). The genome is flanked by 5′ and 3′ untranslated regions (UTRs). The 3′ UTR is polyadenylated, and the 5′ end of the genome is highly structured, containing an internal ribosome entry site (IRES) and a 5′-covalently bound viral protein, VPg (also known as 3B) ^2^. All enteroviruses encode a large polyprotein in a single long open reading frame (ppORF). Human *E. alphacoxsackie*, *E. betacoxsackie*, and around half of *E. coxsackiepol* genomes contain an upstream open reading frame (uORF) that partially overlaps the IRES and the ppORF and encodes a small upstream protein (UP) that promotes virus infection in gut epithelial cells ^5^. Where present, the translation of the uORF produces a peptide of 56–76 amino acid (aa) residues, with a molecular mass of 6.5–9.0 kDa and isoelectric point (pI) of 8.5–11.2^5,6^.

The 5′ UTR contains six stemloop (SL) structures, domains I-VI. Domain I has a role in directing negative-strand RNA synthesis, whereas domains II-VI form a type I IRES, that facilitates cap-independent ribosomal recruitment ^7^. Domain IV is essential in 43S ribosomal preinitiation complex assembly and polyprotein synthesis ^8^. The uORF AUG (uAUG) is situated in domain VI of the IRES (Fig. 1A) ^5^. Mutating the uAUG in the virus context results in a non-viable virus since ppORF translation is entirely IRES-dependent and impossible without the SL-VI uAUG that facilitates initiation complex pre-assembly ^9^. This translational co-dependence complicates investigation of how the expression of these two ORFs (ppORF and uORF) is regulated. Using UP knockout (KO) viruses, ribosome profiling, and IRES-containing reporter systems, we previously demonstrated translation of both ORFs during virus infection, overturning the previous dogma that enteroviruses encode a single polyprotein ^5^.

Here, we identify an interesting feature whereby several enteroviruses contain two upstream AUG codons, both of which are utilized for translation, thus further expanding the translational potential of enteroviruses. We demonstrate the modulation of translation initiation and reading frame usage through the additional upstream AUG. We use several computational and experimental techniques to understand the evolutionary conservation of additional upstream AUGs and their translational potential, regulation, and competitive advantage in terminally differentiated intestinal organoids and neuronal cultures.

## Results

### Analysis of enterovirus sequences reveals additional upstream AUG codons in the vicinity of the IRES SL-VI AUG

A total of 9347 enterovirus sequences were analyzed for the presence of additional upstream AUGs. Using BLASTCLUST, sequences were grouped into 41 clusters based on an 80% identity threshold in the polyprotein amino acid sequence (see Methods). We found over 3000 sequences to have one or more additional AUGs within 20 nucleotides upstream or downstream of the SL-VI uAUG (Fig. 1A, Fig. S1-S3). Most of these additional AUG codons were found upstream rather than downstream of the SL-VI AUG (Fig. 1A), and additional AUGs were present in both rhinoviruses and enteroviruses (Fig. S3). However, the majority of sequences with an additional upstream AUG fell into the “enterovirus A”, “enterovirus C” and “enterovirus D” clusters (Fig. S1), and in fact, these represented the majority of all sequences for the “enterovirus C” and “enterovirus D” clusters (Fig. S3). We define these additional non-SL-VI AUGs as “alternative upstream AUGs”.

In rhinoviruses, the SL-VI AUG is positioned close to the ppORF AUG, whereas in enteroviruses the SL-VI AUG is positioned ≥50 nt and often ≥150 nt upstream of the ppORF AUG (Fig. S3) providing space for a protein-coding uORF. As discussed in Lulla *et al*. (2019) ^5^, a lengthy SL-VI AUG-initiated uORF is present in the majority of sequences in the “enterovirus A” and “enterovirus B” clusters and a substantial number of sequences in the “enterovirus C” cluster, besides some less-abundantly-sequenced clades such as the “enterovirus E”, “enterovirus G” and “porcine enterovirus 9” clusters (Fig. S3). This is reflected in the peak at ∼60–75 codons in a histogram of SL-VI AUG-initiated uORF lengths across all 9347 sequences (Fig. 1B, upper panel). In contrast, in most cases the ORF beginning with an alternative upstream AUG codon (where present) was found to be very short (Fig. 1B, right panel). However, there were some cases where the ORF beginning with the alternative upstream AUG codon had a length of ∼70 codons (Fig. 1B, lower panel). Looking more closely, we found that in 44 and 22 sequences in the “enterovirus C” and “enterovirus A” clusters, respectively, the alternative upstream AUG initiated a uORF with ≥40 codons upstream of the ppORF (Fig. S2, Fig. S3). In some of these sequences (12 “enterovirus C”, 22 “enterovirus A”, and the singleton sequence AF326750), the alternative upstream AUG supplanted the SL-VI AUG as a suitable initiation site for a UP-encoding uORF (according to the Lulla *et al*. 2019 definition ^5^, namely an ORF which overlaps the ppORF by at least 1 nt, is not in-frame with the ppORF, and contains at least 150 nt upstream of the ppAUG). Given these findings, we wanted to investigate whether some enteroviruses could indeed utilize alternative upstream AUG codons for translation initiation.

**Figure 1.**
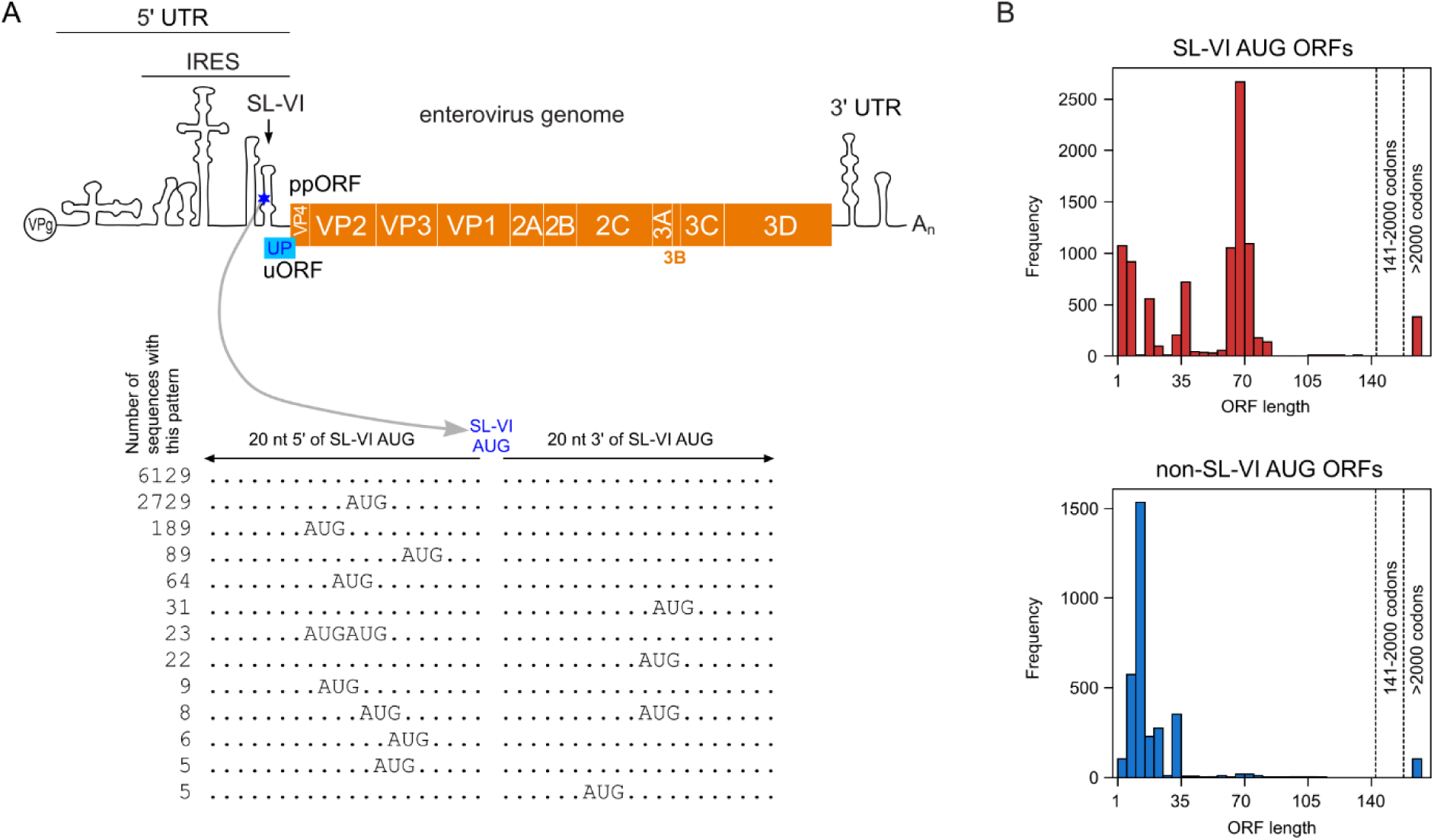
Analysis of AUG triplets in regions flanking the enterovirus SL-VI AUG. (**A**) Schematic representation of the enterovirus genome with indicated features: 5′ and 3′ untranslated regions (UTRs), internal ribosome entry site (IRES), and stemloop VI (SL-VI). Below are the most frequent non-SL-VI AUG patterns in the 20 nt 5′ and 20 nt 3′ of the SL-VI AUG. AUG configurations present in ≥5 of the 9347 sequences analyzed are shown; see Figures S1-S3 for the complete analysis. (**B**) Histograms of SL-VI AUG and non-SL-VI AUG ORF lengths. Histogram bins are in increments of 5 codons from 1–5 up to 136–140; the last two bins are for lengths 141–2000 and >2000 codons; ORF lengths in the last bin correspond to cases where the upstream-AUG-initiated ORF is contiguous with the ppORF (often rhinoviruses). ORF lengths are counted for each relevant AUG even if there are multiple in-frame AUGs.

### Analysis of *E. alphacoxsackie* and *E. coxsackiepol* clades that encode a UP-like protein initiated at an alternative upstream AUG codon

Looking first at the 22 *E. alphacoxsackie* sequences, besides the singleton sequence AF326750, we found the additional AUG (uuAUG, upstream upstream AUG) to be located 10 nt upstream of the SL-VI uAUG. Translation from the uuAUG in these sequences would yield a protein closely related to the characterized UP ^5^. Aside from an altered N-terminus (MVTI in the uORF compared to MAAY in the uuORF), the predicted protein possesses typical UP features, including a size of 8–9 kDa, a transmembrane helix (TMH) domain ^10^, a conserved WIGHP sequence, and a high isoelectric point (9.5–10.7) (Fig. 2A). The canonical uAUG-initiated short ORF in these isolates was truncated by an in-frame stop codon at the tip of SL-VI, resulting in a 4 aa peptide upon translation (Fig. 2B). An analysis of metadata associated with these sequences revealed that the viruses were collected from different geographical locations (India, Bangladesh, China, Cameroon, Cambodia, and the Netherlands), mostly isolated from gastrointestinal samples, and often associated with paralysis or myelitis (Table S1).

**Figure 2.**
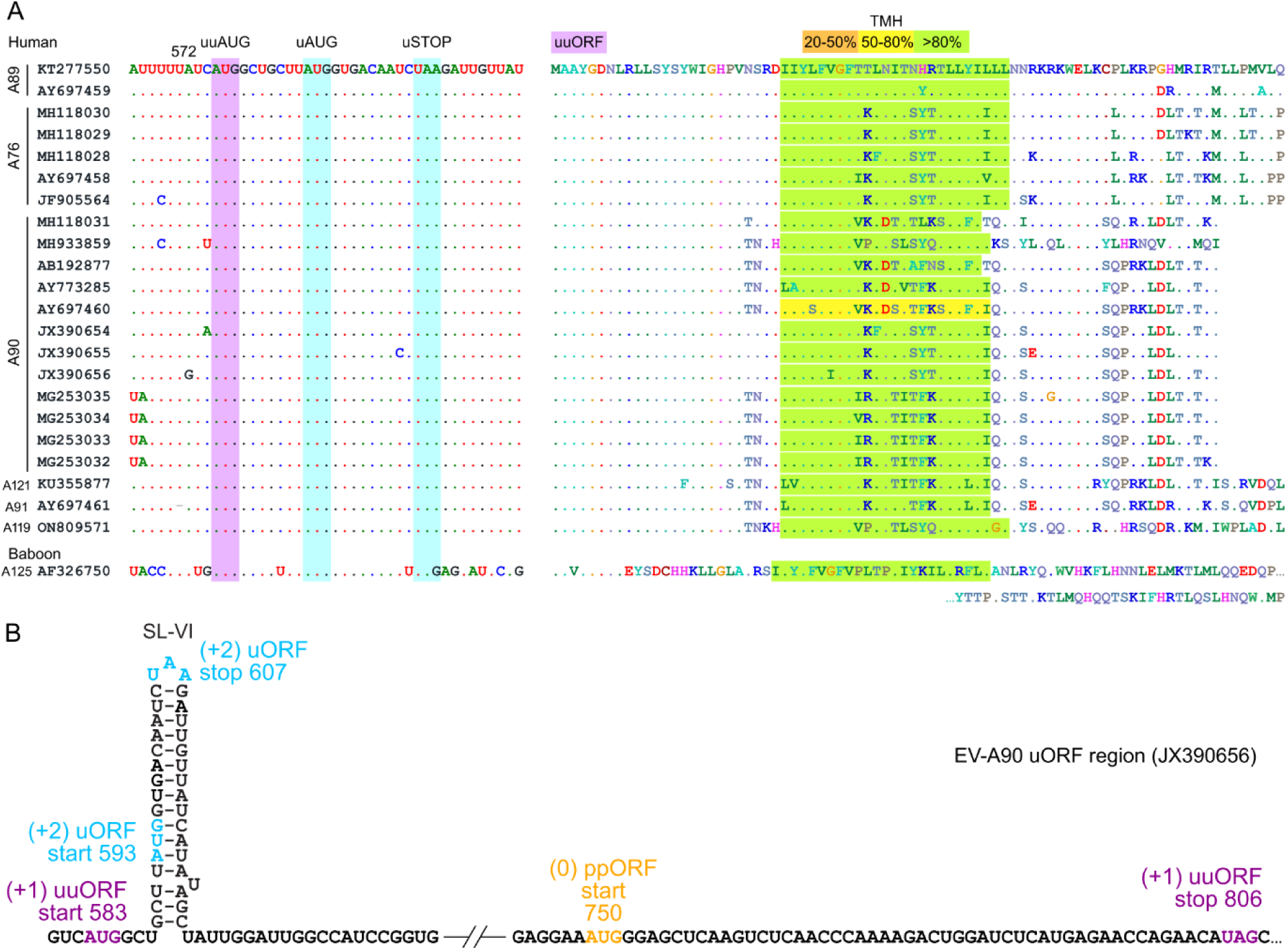
*E. alphacoxsackie* sequences that encode a UP-like protein initiated at an alternative upstream AUG codon. (**A**) Nucleotide (left) and uuORF-encoded protein (right) sequences in enterovirus A89/A76/A90/A121/A91, where the uAUG-initiated uORF is truncated and the uuAUG-initiated uuORF can potentially rescue UP expression. Transmembrane helix (TMH) predictions are highlighted in yellow (50–80% confidence) or green (>80% confidence). Two ORFs are indicated in purple (uuORF) and blue (uORF) highlighting. (**B**) Schematic representation of the EV-A90 (JX390656) IRES dVI region with uORF (blue), uuORF (purple), and ppORF (orange) start and stop codons annotated.

The singleton sequence AF326750 (Fig. 2A) originated from a baboon stool sample ^11^. Its UP sequence and properties were divergent from the other sequences, with a much longer protein (with predicted TMH) and the absence of the conserved WIGHP sequence (Fig. 2A).

Looking at the 12 *E. coxsackiepol* sequences, we also found the additional AUG (uuAUG) to be located 10 nt upstream of the SL-VI uAUG. Translation from the uuAUG in these sequences would yield a protein closely related to the characterized UP ^5^. Aside from an altered N-terminus, the predicted protein meets the above-mentioned criteria, albeit with fewer sequences having a predicted TMH (Fig. 3). The canonical uAUG ORF in these isolates is truncated by an in-frame stop codon, resulting in a 4–18 aa peptide upon translation (Fig. 3).

The existence of a double AUG signature in these two different groups of *Enterovirus* species provides additional capacity for coding if both AUGs are translationally competent. This may be particularly important for those enterovirus strains where a long uORF is initiated from the uuAUG instead of the uAUG (Fig. 2, Fig. 3). Taken together, these analyses suggest alternative UP-coding possibilities, which have been previously overlooked.

**Figure 3.**
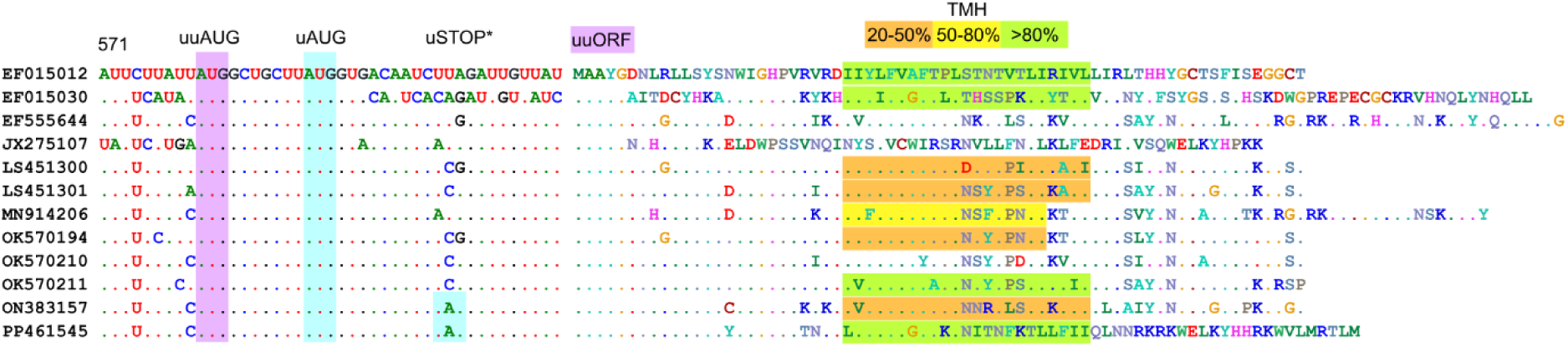
*E. coxsackiepol* sequences that potentially encode a UP-like protein from an alternative upstream AUG codon. Nucleotide (left) and uuORF-encoded protein (right) sequences in enterovirus CVA-19/CVA-1/C113/C116/CVA-22, where the uORF is truncated and the uuORF can potentially rescue UP expression. Transmembrane helix (TMH) predictions are highlighted in orange (20–50% confidence), yellow (50–80% confidence) or green (>80% confidence). Two ORFs are highlighted in purple (uuORF) and blue (uORF). uSTOP* indicates stop codons in the uORF for the last two sequences; the remaining ten have stop codons further downstream, resulting in a 17–18 aa peptide.

### Both upstream AUGs can be used for translation initiation in enterovirus CVA-13

To test the possibility of translation initiation at uAUG and uuAUG codons during virus infection, we utilized ribosome profiling (Ribo-Seq), a technique for the global footprinting of translating ribosomes. To identify translation initiation sites, we used the translation inhibitor lactimidomycin (LTM), which acts on initiating ribosomes ^12^. This approach has proven valuable to detect closely located translation initiation events during RNA virus infection ^13^. We used the CVA-13 Flores strain of *E. coxsackiepol* that contains both the SL-VI uAUG and a uuAUG 10 nt upstream, with the uAUG-initiated ORF coding for the UP (Fig. 4A). The infection dynamics of this virus followed a classical enterovirus scenario as previously observed for *E. alphacoxsackie*, *E. betacoxsackie* and *E. coxsackiepol* strains ^5^, so we selected two representative time points corresponding to early (5 hpi) and late (7 hpi) infection (Fig. 4B). Profiling of initiating ribosomes revealed two distinct initiation peaks corresponding exactly to the uuAUG and the uAUG (Fig. 4C), indicating that translation initiation can occur at either AUG. We conducted a similar experiment without LTM pre-treatment to confirm that this was not an artifact of antibiotic treatment. Interestingly, both the uuORF and the uORF were occupied by translating ribosomes, despite the uuORF only coding for an 8 aa peptide (Fig. 4D).

Surprisingly, the uORF and uuORF initiation peaks were higher than the ppORF initiation peak in the LTM-treated samples (Fig. 4C), contrary to the ratio of expression levels expected from previous work ^5^ and subsequent functional analysis (see below). This could potentially result from ligation biases or an artifact of LTM treatment ^14^. To investigate the latter possibility, we selected cellular mRNAs with upstream AUGs and compared ribosome occupancy levels on the uAUGs and the main ORF AUGs (mAUGs) (Fig. S4). Due to late infection-associated host shut-off, the 7 hpi LTM-treated libraries had very few host-mapping reads. In the 5 hpi LTM-treated libraries, the mean host uAUG:mAUG occupancy ratio was ∼0.31 whereas for the virus this ratio was ∼10.8. The analysis comes with many caveats: relatively few host mRNAs with suitable uAUGs, differences in uAUG-mAUG spacing and other 5′ UTR features between host and viral mRNAs, unquantified levels of alternative splice forms, different ribosome loading levels of host and viral transcripts, etc (see Methods). However, the analysis of host mRNAs does indicate that the high uAUG:mAUG ratio seen for the virus (∼35-fold difference from host) is likely not simply an artifact of LTM treatment, but might instead be an IRES-specific effect of the drug whereby the magnitudes of the uORF and uuORF initiation peaks do not reflect their translation levels.

We assessed Ribo-Seq quality by plotting histograms of ribosome-protected fragment (RFP) mapping positions relative to annotated initiation and termination sites in host mRNAs (Fig. S5A-B), and length and triplet phasing distributions of RFPs mapping to virus or host mRNA coding regions (Fig. S5C-D). The read length distribution of virus-mapping reads was sharply peaked at 27–29 nt (Fig. S5C). However, the read-length distribution for the host-mapping reads was less sharply peaked, especially at the later time point (7 hpi). As explained in Lulla *et al*., (2019) ^5^, we take this to be a consequence of severe virus-induced shut-off of host translation which, for the host, greatly reduces the proportion of *bona fide* RPFs to contamination at 7 hpi, hence leading to a host-specific reduction in Ribo-Seq quality. To maximize the contribution of *bona fide* RPFs for both the host and the virus, we restricted all subsequent analyses to 27–29 nt reads only. The majority of virus-mapping read 5′ ends mapped to the first nucleotide of codons (Fig. S5D), indicating that the Ribo-Seq datasets were overall of high quality. For the host, even after restricting to 27–29 nt reads, the phasing quality was noticeably degraded (more reads mapping to the 2nd and 3rd positions of codons), especially at 7 hpi. Analysis of host mapping reads confirmed that LTM-treated samples mapped initiating ribosomes (Fig. S5A) and non-treated samples mapped elongating ribosomes (Fig. S4B). The full-virus-genome translation initiation (Fig. S5E) and elongation (Fig. S5F) profiles were consistent with previously analyzed enterovirus Ribo-Seq datasets ^5,13^. The noisier LTM virus profile at 7 hpi (Fig. S5E) could be linked to a restricted ability of apoptotic cells to take up the drug.

Therefore, ribosome profiling experiments confirm that both upstream AUGs can be used for translation initiation during enterovirus CVA-13 infection. The relative usage of these ORFs can be defined using reporter systems ^5^.

**Figure 4.**
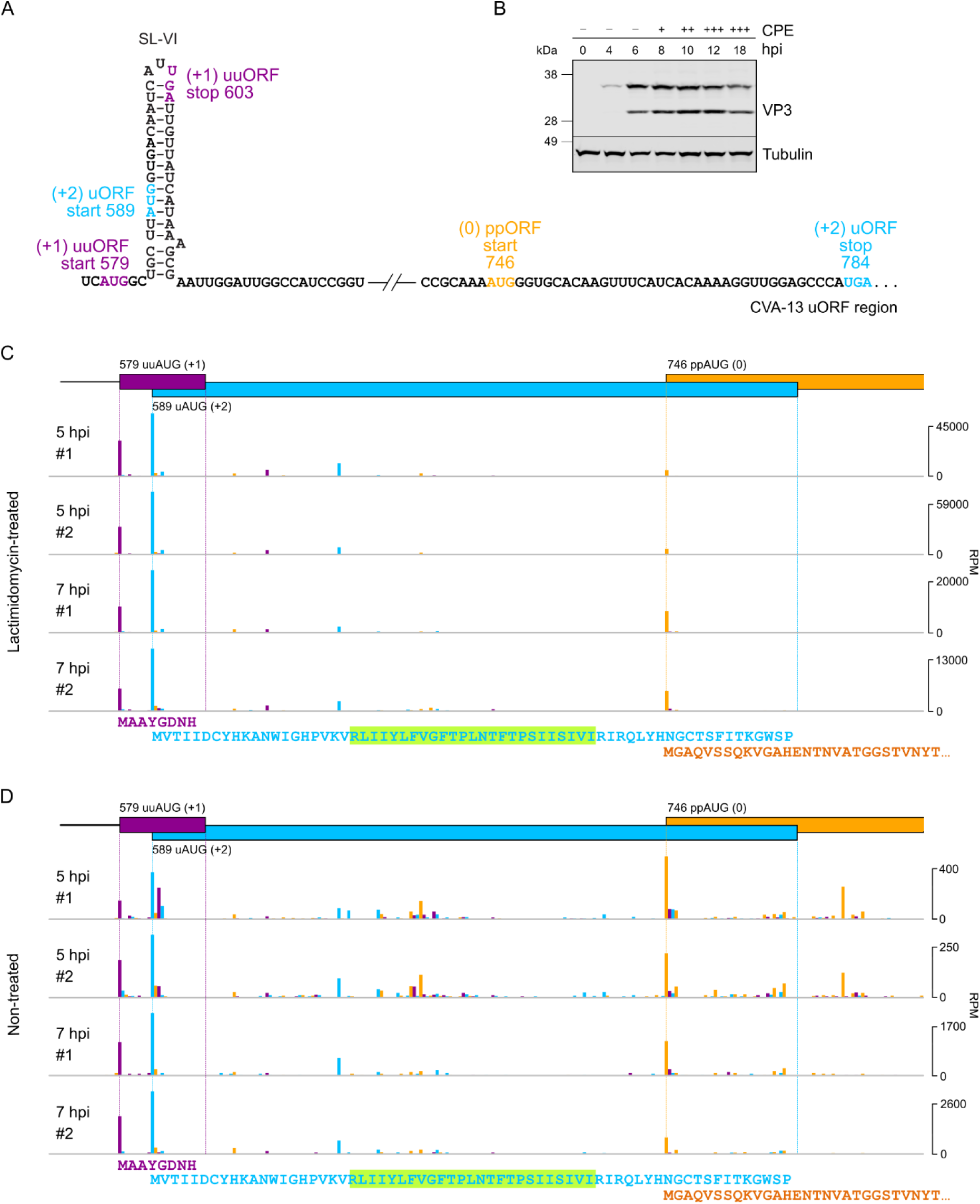
Translation initiation in enterovirus CVA-13. (**A**) Schematic representation of the CVA-13 IRES dVI region with the uORF (blue), uuORF (purple) and ppORF (orange) start and stop codons annotated. (**B**) Analysis of viral protein expression in HeLa cells infected with CVA-13. Cells were infected at an MOI of 10, harvested at 0–18 hpi as indicated, and accumulated virus structural protein VP3 was analyzed by western blotting with the anti-VP3 antibody. Observed cytopathic effect (CPE) of virus infection is indicated by (−) absence, (+) <40% CPE, (++) 40–80% CPE, (+++) >80% CPE. (**C**) Ribosome profiling of CVA-13-infected cells at 5 and 7 hpi in the presence of translation initiation inhibitor lactimidomycin. (**D**) Ribosome profiling of CVA-13-infected cells at 5 and 7 hpi without lactimidomycin treatment. (C-D) Ribo-Seq RPF densities (mapping positions of 5′ ends of reads with a +12 nt offset to indicate the approximate P site) in reads per million mapped reads (RPM). Colors indicate the triplet phase of reads relative to the genome start. Since most read 5′ ends map to the first nucleotide of codons (Fig. S5D), reads deriving from translation in the uuORF (+1), uORF (+2) and ppORF (0) are expected to be predominantly purple, blue and orange respectively. The amino acid sequences of the predicted proteins/peptides encoded by the three ORFs are displayed underneath the profiles. The highlighted green region indicates the predicted transmembrane domain of the UP protein encoded by the uORF. Full-genome plots and Ribo-Seq quality control plots are provided in Fig. S5.

### Evaluation of long uORF and ppORF translation using an IRES reporter system

We previously observed that ribosome profiling may not provide accurate information on relative translation efficiencies for short ORFs due to ribosome pausing and Ribo-Seq library preparation biases ^5,13,15^. Therefore, to evaluate the translation efficiency of long uORFs and the ppORF in different representative enterovirus species (Fig. 2-3), we employed a previously developed dual-luciferase reporter system ^5^, where a 2A-FFLuc cassette was placed in the 0, +1 or +2 frame relative to the ppORF (Fig. 5A). We chose three different representative virus strains, with differing uuORF/uORF configurations: a UP-coding uORF in the +2 frame with a truncated uuORF in the +1 frame (CVA-13); a UP-like-coding uuORF in the +2 frame with a truncated uORF in the 0 frame (CVA-1); and a UP-coding uuORF in the +1 frame with a truncated uORF in the +2 frame (EV-A90). The CVA-13 sequence was derived from the clinical isolate used for ribosome profiling (Fig. 4), whereas the 5′ UTRs of CVA-1 and EV-A90 were synthesized *in vitro*. The CVA-1 IRES (JX174177) was derived from the virus isolated from the stool sample (Fig. S6), and the EV-A90 IRES (JX390656) was derived from a sequence from an acute flaccid paralysis patient (Fig. 2). Consistent with the uORF translation levels seen previously in echovirus 7 (4–5% relative to the ppORF) ^5^, expression of the longer upstream ORF in the three new enterovirus cassettes was in the range 3–6%, depending on cell line and virus strain (Fig. 5). Translation initiation at the AUG codon for the shorter (truncated) upstream ORF would lead to early termination, upstream of the firefly luciferase reporter. Therefore, for the remaining constructs, firefly activity measures background – likely including low-level initiation at various non-AUG sites upstream of the firefly coding sequence. These background levels were in the range 1–3%. For all three virus strains, translation of the longer upstream ORF was significantly higher than the background level (Fig. 5), thus confirming that UP can be translated from either a uAUG or a uuAUG.

**Figure 5.**
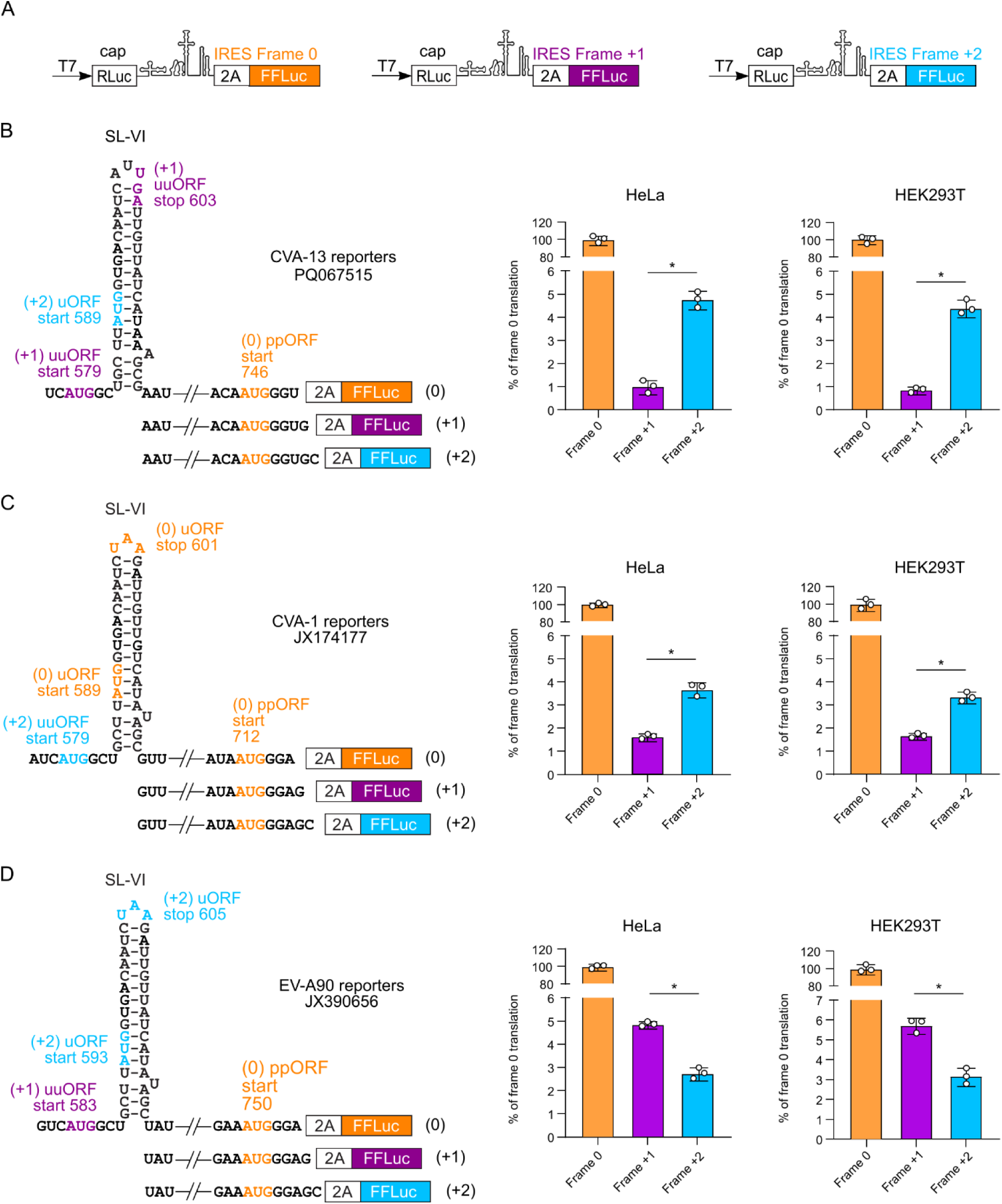
Usage of both upstream AUGs in different enterovirus IRES reporters. (**A**) Schematic representation of the modified pSGDluc expression vectors used to measure translation at the polyprotein (ppORF, orange) and two upstream (blue and purple) AUG codons. (**B**-**D**) Analysis of IRES activities for ppORF, uORF and uuORF expression relative to cap-dependent expression by dual-luciferase reporter assay in HeLa and HEK293T cells (FFLuc/RLuc) at 8 h post-transfection for CVA-13 (B), CVA-1 (C) and EV-A90 (D) enterovirus IRESes. IRES activities in each of the three frames were normalized to cap-dependent signal and presented as relative activities in the three frames (means ± SD, *n* = 3 biologically independent experiments). Each panel contains a schematic representation of the corresponding IRES reporters (left). Statistical analysis of the difference between the translation efficiency of frames +1 and +2 was conducted using two-tailed *t*-tests; * *p* value ≤ 0.05.

### Functional significance of the uuAUG in enterovirus CVA-13

To address the functional significance of a non-UP-encoding uuAUG in the virus context, the enterovirus CVA-13 Flores strain (utilized for ribosome profiling, Fig. 4) was used to develop a CVA-13 reverse genetics construct. This was made by fusing the entire CVA-13 genome sequence after a T7 promoter (Fig. 6A). We confirmed that the recombinant CVA-13 virus was stable on passaging and demonstrated growth properties similar to the parental clinical isolate (Fig. 6B). A CVA-13-uuKO recombinant virus was created with an AUG→GUG substitution to the uuAUG (Fig. 6A). This substitution was already present in minor quantities in the clinical CVA-13 isolate (Fig. S7) and is predicted to knock down translation of the 8 aa uuORF peptide while preserving the translation of both other frames (uORF and ppORF). We confirmed the expression of UP in all three viruses, suggesting the uuAUG-independent translation of the two other frames (Fig. 6C). We used several alternative approaches to probe the functional consequences of preventing uuORF expression. First, the WT and CVA-13-uuKO viruses were competed in the highly susceptible HeLa cell line. Both viruses were retained in the population for six passages in four independent experiments, confirming no advantage/disadvantage of the uuKO mutation in HeLa cells (Fig. 6D).

**Figure 6.**
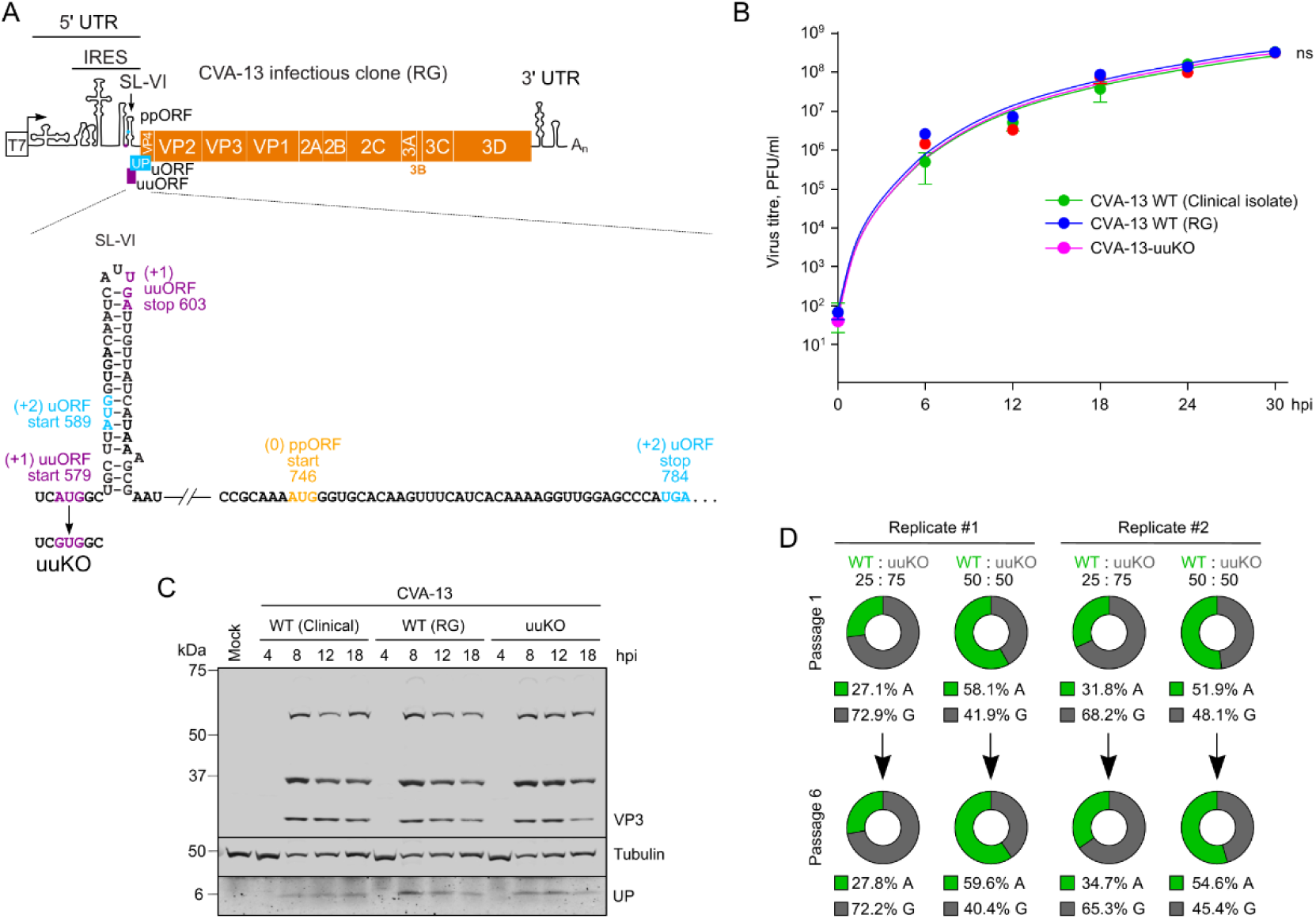
Development of the reverse genetics (RG) plasmid for CVA-13 and properties of the uuORF knockout virus in HeLa cells. (**A**) Schematic representation of the CVA-13 infectious clone with the zoomed-in SL-VI/uORF region indicating the uuKO mutation (AUG→GUG). (**B**) Multistep growth curves of CVA-13 viruses in HeLa cells (MOI 0.1). (**C**) Analysis of VP3 and UP expression in CVA-13-infected HeLa cells. Cells were infected at MOI 10, harvested at the indicated time post-infection, and analyzed by western blotting. (**D**) CVA-13 WT and uuKO virus competition was performed in HeLa cells. Cells were infected with WT and uuKO mutant viruses at the indicated proportions in four independent experiments and passaged six times at MOI 0.01. The virus RNA from passages 1 and 6 was isolated, RT-PCR amplified, and sequenced. The proportion of A^579^(WT)/G^579^(uuKO) nucleotide abundance was determined for each sample.

We then hypothesized that presence of the uuAUG might be important in terminally differentiated cells, such as intestinal epithelia or neurons – natural infection sites for many enteroviruses ^4^. Therefore we developed an infection system for CVA-13 in differentiated iPSC-derived i^3^Neurons ^16,17^ and assessed the replication kinetics of WT and uuKO viruses. We confirmed efficient virus replication in this system but did not observe any differences between the growth properties of the tested viruses (Fig. 7A). However, a competition experiment using two different WT[AUG]:uuKO[GUG] ratios (30:70 and 70:30) revealed a shift towards WT virus in the infection mixture when the WT:uuKO ratio was 30:70 (Fig. 7B), suggesting an advantage of the uuAUG-containing virus (WT) in this system.

An advantage of WT over uuKO CVA-13 was also observed in differentiated human intestinal organoids. The growth of individual viruses was nearly identical for all three viruses (Fig. 7C and Fig. S7), resulting in productive virus release without any cytopathic effect, suggesting possible persistent infection of CVA-13. Due to the limited lifespan of differentiated organoid cultures (4-5 days post-differentiation), assessing this in longer-term conditions was not possible. Interestingly, the more sensitive competition experiment using two different WT[AUG]:uuKO[GUG] ratios (30:70 and 70:30) revealed a shift towards WT virus in the mixture when the WT:uuKO ratio was 30:70 (Fig. 7D), recapitulating the results observed in differentiated neurons and confirming an advantage of the uuAUG-containing virus (WT) in human intestinal organoids. No changes were detected in either system when the initial WT:uuKO ratio was set at 70:30, suggesting no competitive pressure when WT is already over-represented in the mixture (Fig. 7B and Fig. 7D).

Taken together, our results suggest that mutation of the uuAUG has no effect in the HeLa cell line but resulted in an apparent phenotype in competition assays using physiologically relevant systems: terminally differentiated neurons and intestinal organoids. These results echoed the UP phenotype, which was only evident in the organoid system ^5^, and emphasize the role of IRES elements in differentiated cells/tissues. To investigate the mechanistic role of the uuORF, we decided to assess other elements of the uuORF in a sensitive reporter system.

**Figure 7.**
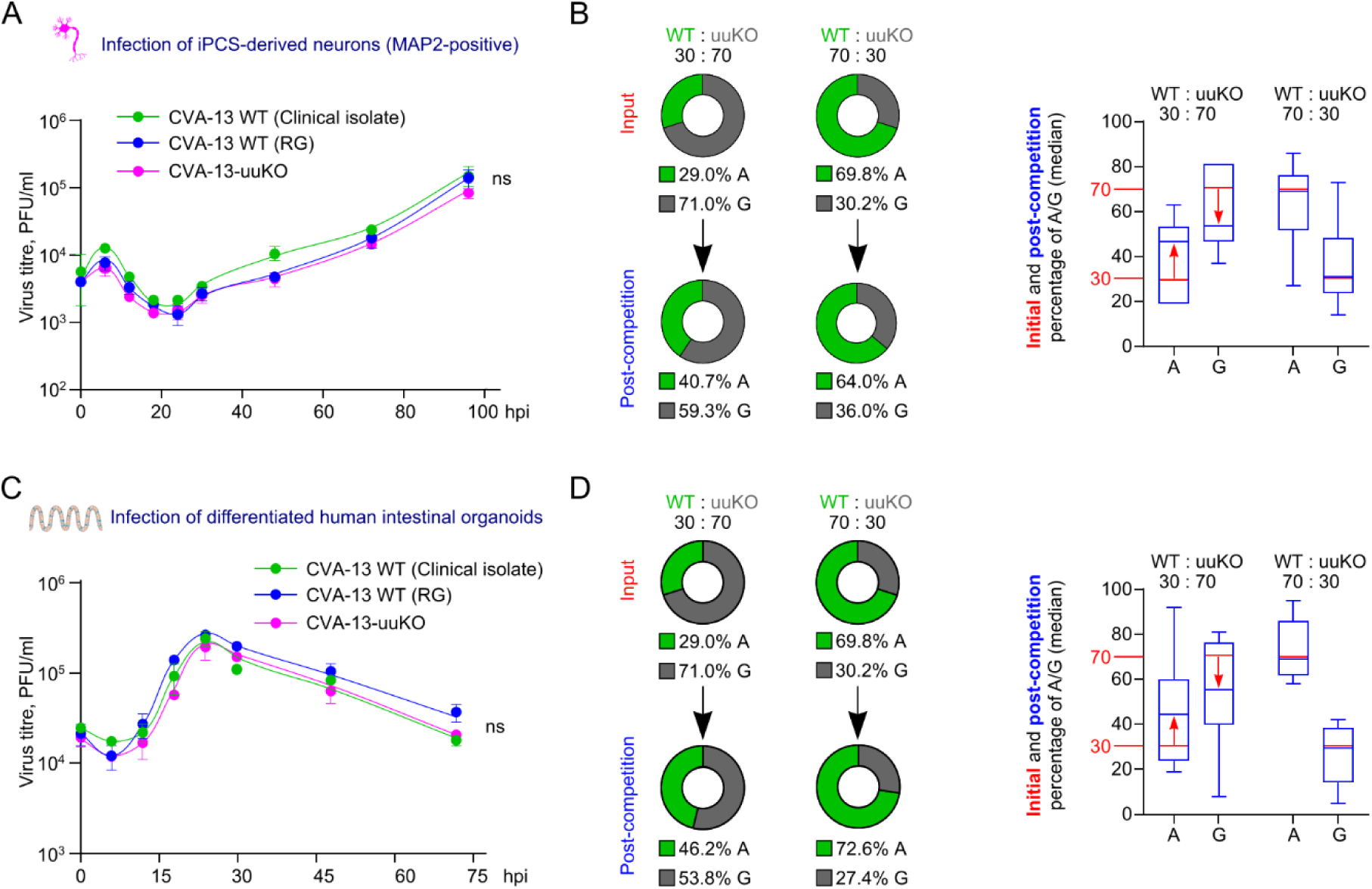
Properties of the uuORF knockout virus in the neuronal and intestinal organoid infection models. (**A**) Differentiated iPSC-derived neurons were infected in quadruplicate at MOI 0.1, and the released virus was analyzed by plaque assay in HeLa cells. (**B**) CVA-13 WT and uuKO virus competition was performed in differentiated iPSC-derived neurons. Neurons were infected at MOI 0.1 with WT and uuKO mutant viruses at the indicated proportions in 6 independent experiments and incubated for 6 days with media changes every other day. The virus RNA from input and post-competition samples was isolated, RT-PCR amplified, and sequenced. The proportion of A^579^(WT)/G^579^(uuKO) nucleotide abundance was determined for each sample and presented as a doughnut plot (left, mean values) and box and whiskers plot with boxes extending from the 25^th^ to 75^th^ percentiles, and whiskers ranging from the smallest to largest values (right, median values). (**C**) Monolayers of differentiated cultured intestinal organoids were infected in quadruplicate with CVA-13 viruses at MOI 10, aliquots of culture media were collected at the indicated time points, and viral titers were analyzed by plaque assay in HeLa cells. (**D**) CVA-13 WT and uuKO virus competition was performed in differentiated human intestinal organoids. Organoid monolayers were infected at MOI 1 with WT and uuKO mutant viruses at the indicated proportions in 10 independent experiments and incubated for 4 days with media changes after 24 and 48 hpi. The virus RNA from input and post-competition (96 hpi) samples was isolated, RT-PCR amplified, and sequenced. The proportion of A^579^(WT)/G^579^(uuKO) nucleotide abundance was determined for each sample and presented as a doughnut plot (left, mean values) and box and whiskers plot with boxes extending from the 25^th^ to 75^th^ percentiles, and whiskers ranging from the smallest to largest values (right, median values). The data in (A) and (C) represent two biologically independent experiments; ns, nonsignificant using nonlinear regression analysis. The sequencing of the RT-PCR product derived from virus collected at 96 hpi (neuron) and 72 hpi (organoid) is provided in Fig. S7.

### A role of the CVA-13 uuORF in translation regulation of IRES-dependent ORFs

To investigate a potential role for the uuAUG in modulating translation of other ORFs, we utilized the above-mentioned CVA-13 reporter system. However, we aimed to improve it by including the viral 3′ UTR as it was previously shown that IRES function and its cell type-specific restrictions may be influenced by *cis*-acting interactions between the IRES and the Z domain of the 3′ UTR ^18,19^. In addition, we optimized the reporter assay in human intestinal epithelial cells (HIEC6) as a more relevant model system to evaluate enterovirus-specific translation. The uuKO mutation was introduced in both versions of the reporter construct (with and without 3′ UTR) and for firefly luciferase in each of the three frames (Fig. 8A). Interestingly, the relative usage of the functional ppORF (0) and uORF (+2) frames did not change between the original and 3′ UTR-containing reporters; however, the overall translation was increased 5–6 fold (Fig. 8B), confirming the importance of the 3′ UTR in IRES-dependent translation. We used both reporters to evaluate the effect of the uuKO mutation in the reporter systems. In agreement with assays in infected HeLa cells (Fig. 6), in most tested conditions, we did not observe any differences between WT and uuKO reporters in either cell line (Fig. 8C-D, Fig. S8). We also added viral RNA to transfection mixtures to mimic the infection condition since viral proteins such as 2A were previously shown to affect IRES- and cap-dependent translation ^20^. Overall, adding virus RNA also did not result in differences between WT and uuKO reporters (Fig. 8C-D, Fig. S8).

Although we did not observe any effect upon mutation of the uuAUG, we also wanted to test whether the position of the uuORF stop codon might affect translation of the other ORFs. The positions of stop codons in short uORFs/uuORFs vary between viruses. Out of 9333 analyzed sequences (Fig. S9), 3975 have a short AUG-to-stop-codon ORF in any frame (where short ORF here means that the AUG is anywhere up to 20 nt 5′ of the SL-VI AUG, and the stop codon is anywhere up to 30 nt 3′ of the SL-VI AUG). Of these, 1545 have a stop codon in-frame with the SL-VI AUG within the 18 nt 3′ of the SL-VI AUG, i.e. they have a SL-VI AUG ORF of less than or equal to 6 sense codons (Fig. S9).

We hypothesized that removing stop codon(s) to turn short ORFs into longer ORFs might affect translation in the other IRES-initiated ORFs. In some viruses, there are two stop codons in SL-VI in the same frame. For example, in CVA-13 the second stop codon is situated directly opposite the conserved SL-VI uAUG (Fig. 8C), making it difficult to manipulate without breaking the SL-VI structure and IRES function ^5,9,21^. However, for the two previously used IRES examples (Fig. 5C-D), the short ORF frame has a stop codon that is located in the loop of SL-VI. Therefore, these SL-VI stop codons could be manipulated, similar to a previously used uORF knockout strategy (Loop mutant) ^5^. In the first example – the CVA-1 IRES – the stop codon for the short SL-VI uAUG-initiated ORF is in the same frame as the ppORF and there are also two additional stop codons present in this frame in the linker region at positions 640 and 661 (Fig 8E). Mutation of the (0) frame SL-VI stop to an isoleucine codon (AUU) affected the relative expression levels in the (+2) frame (Fig. 8E). In the second example – the EV-A90 IRES – mutation of the SL-VI stop codon decreased the (+1) uuORF expression (Fig. 8F), suggesting that the upstream ORF expression can be modulated through the SL-VI stop codon. Additional stop codons are present in the (+2) frame downstream of the EV-A90 SL-VI (positions 644, 689 and 731). Taken together, these results suggest a modulatory role of the stop codon in SL-VI.

**Figure 8.**
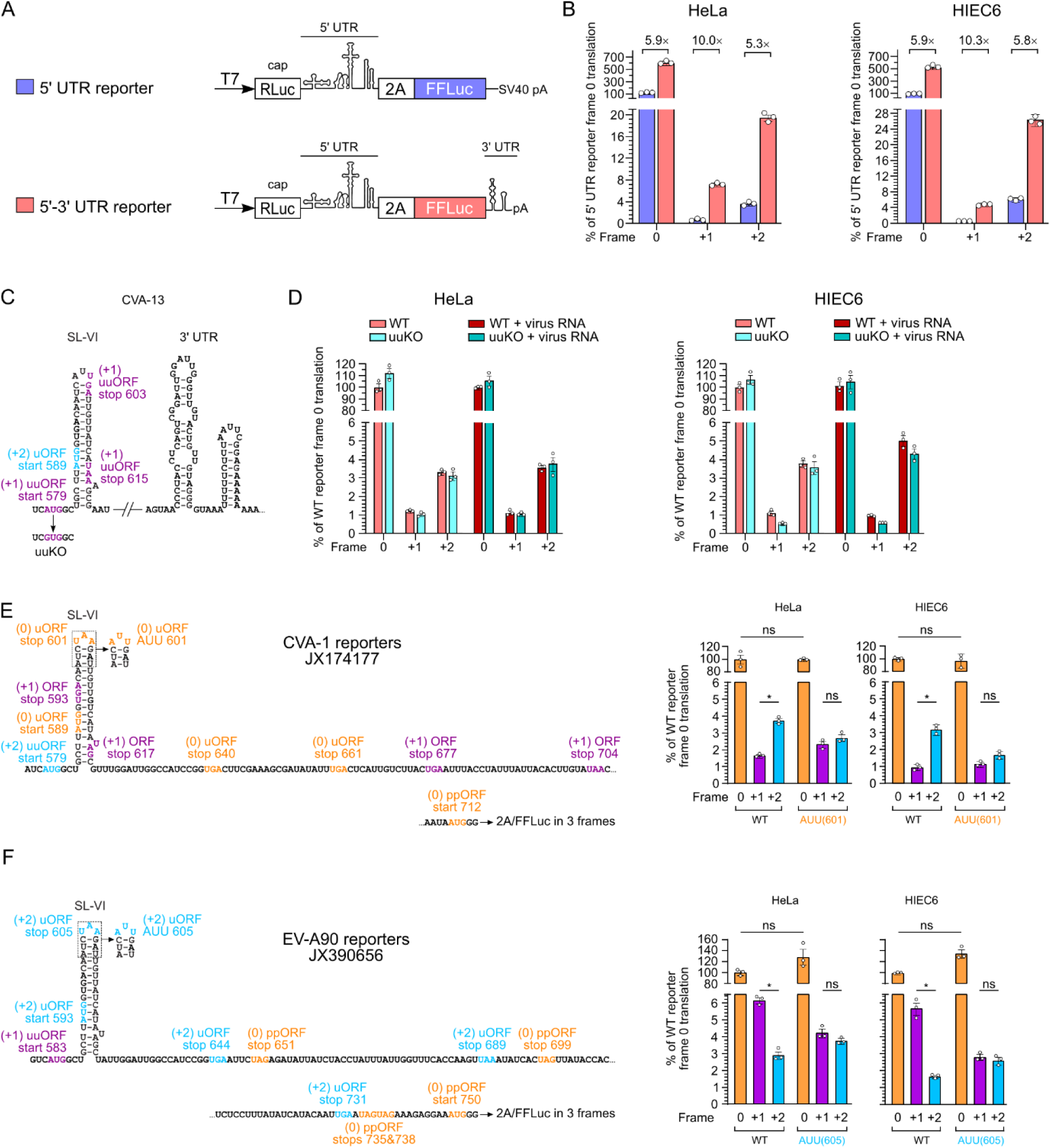
The role of uuORF start and stop codons in IRES-dependent translation. (**A**) Schematic representation of the 5′ UTR and 5′-3′ UTR reporters used to measure translation in the three frames. (**B**) Analysis of IRES activities for CVA-13 reporters in the three frames in the 5′ UTR and 5′-3′ UTR reporters in HeLa and intestinal HIEC6 cells. Fold changes between reporter activities are indicated above each pair. (**C**) Schematic representation of the SL-VI domain of the WT and uuKO mutant CVA-13 sequence and the 3′ UTR translation-enhancing element. (**D**) Analysis of IRES activities for CVA-13 5′-3′ UTR reporters in the three frames in HeLa and intestinal HIEC6 cells with and without virus RNA at 8 h.p.t. The full set of 5′ UTR and 5′-3′ UTR reporter data can be found in Fig. S8. (**E-F**) Schematic representation of the CVA-1 (E) and EV-A90 (F) reporters and their AUU mutants used to measure translation in three frames in HeLa and HIEC6 cells. Statistical analysis was conducted using two-tailed *t*-tests; * *p* value ≤ 0.05; ns, non significant.

## Discussion

Here, we show that many enteroviruses have two upstream AUG codons, the SL-VI uAUG and an additional AUG (uuAUG). Combining several computational and experimental approaches, we address the significance of the additional upstream AUG in enterovirus genomes. First, using ribosome profiling of CVA-13-infected cells, we confirm that both upstream AUG codons can be used for translation initiation (Fig. 4).

Regulation via uORF expression is important in cell/tissue-specific translation, particularly in neural tissues ^22,23^. Enterovirus infections can lead to CNS complications, such as meningitis, encephalitis, acute flaccid paralysis, and other manifestations. The majority of neurovirulence determinants in enteroviruses have been mapped to IRES elements (reviewed in Lulla and Sridhar, 2024) ^4^, highlighting the likely role of translation regulation ^24^. In addition, IRES elements, particularly uAUGs, are often associated with neurovirulence in mice ^25,26^, and IRES-related neurotropic properties have been recently described for the mouse-adapted EV-A71 strain ^27^. Therefore, one possible explanation for a role of the additional upstream AUG in enteroviruses could be tissue-specific regulation of translation. In line with these predictions, we observe that WT (double-uAUG) CVA-13 outcompeted the uuKO virus in both neuronal and intestinal organoid infection models (Fig. 7). These findings suggest that the uuORF in CVA-13 can modulate translation and/or replication in terminally differentiated infection sites, such as intestinal epithelia and neurons.

Upstream ORFs have been characterized for several viral and eukaryotic mRNAs ^5,28–31^. The functional significance and regulatory mechanisms may differ; however, some parallels can be drawn. In coronaviruses, the role of uORFs has been characterized as regulatory but not essential ^32^. Interestingly, in line with our results, Wu *et al.* showed that disruption of a coronavirus uORF enhanced translation of the main ORF *in vitro*, without detected effects on coronavirus replication. The authors also highlight the regulatory role of the uORF and its potential role in long-term survival during persistent infection ^32^. By using a system similar to our bicistronic reporters, it was shown that a uORF modulates L mRNA (polymerase) translation in ebolaviruses in response to cellular stress. In contrast to enteroviruses and coronaviruses, knockout of the ebolavirus uORF led to a one-log decrease in virus titer in several cell lines ^33^, suggesting stronger dependence on the uORF in this virus. The HIV-1 genome encodes a minimal uORF, consisting of a start and stop codon only, overlapping with the Vpu start site. Mutation of either of these two codons results in a five-fold reduction in Env protein expression ^34^, further highlighting short ORF-mediated modulation of translation in different virus families.

We further corroborated our findings using several IRES reporter systems (Fig. 5 and Fig. 8). Interestingly, the additional upstream AUG alone did not provide any advantage to CVA-13 (*E. coxsackiepol*) during infection of susceptible cell lines (HeLa, HIEC6) and when tested in reporter systems. Due to the overlapping nature of functional elements in RNA viruses ^35^, it was not technically possible to address all scenarios in the context of virus infection and/or IRES reporter systems. Nevertheless, we combined several possible approaches and reporter systems derived from three different enteroviruses and found that a truncated uORF affects translation of other ORFs. Interestingly, removing SL-VI stop codons led to dysregulation of long uuORF usage (Fig. 8E-F). There are several possible interpretations of these results. It is possible that the short uORF recruits additional ribosomes, which, after translating the short SL-VI ORF, can then be recycled to translate the long uuORF. When the loop stop codon of the truncated uORF is mutated to AUU, translating ribosomes terminate too far downstream to be recycled back to the uuORF AUG codon, so we observe reduced translation in the (+2) frame (Fig. 8E) and the (+1) frame (Fig. 8F). In analogy to these results, stressed eukaryotic cells can regulate their gene expression via reinitiation after uORF translation ^36^. Alternatively, this effect can be IRES context-specific or cell type-specific (Fig 7). For example, the AUU mutant may alter IRES structure and/or IRES trans-activating factor (ITAF) recruitment and subsequently affect the uuORF to uORF initiation ratio. Interestingly, we recently demonstrated that translation of Zika virus uORFs can modulate main ORF expression, further expanding the uORF-dependent translational modulation ^31^.

In response to stress, cellular uORFs – such as in the mRNA that encodes activation transcription factor 4 (ATF4) – can act as enhancers of main ORF translation ^37^. Upon enterovirus infection, host translational shut-off is a well-established process caused by viral protease 2A-driven cleavage of eIF4G ^38^, a scaffolding initiation factor that is not required for IRES-dependent translation ^39^. It is plausible that additional short uORFs in enteroviruses can also contribute in responding to stress and/or regulation of IRES-dependent translation ^40,41^.

This study further highlights the consideration of multiple upstream AUGs and upstream ORFs when annotating enterovirus genomes. The ability of enteroviruses to encode the UP protein via multiple routes further expands their coding capacities and provides a platform for further investigations into IRES-dependent translation. Furthermore, the cell type-specific preferences demonstrated in this work provide a foundation for further investigations into IRES-mediated pathogenicity and neuro-attenuation mechanisms.

## Materials and Methods

### Bioinformatic analysis of uORF and uuORF characteristics in enterovirus sequences

Enterovirus sequences were retrieved from the National Center for Biotechnology Information (NCBI) nucleotide database on 22 March 2024 by searching with the parameters ’txid12059[Organism:exp] AND 5000:50000[Sequence Length]’, which yielded 11,791 sequences, of which 30 were NCBI RefSeqs. Sequences with PAT (patent), SYN (synthetic) or CON (constructed) labels in the GenBank Division field were removed, leaving 11,469 sequences with the “VRL” (viral) label and 228 sequences with the ENV (environmental sample) label. Sequence accession OP414044 was discarded due to the first ∼1440 nt unexpectedly mostly comprising a perfect inverted repeat of itself. Sequence accessions AB469183 and LC637981 were discarded as, although labelled as “VRL”, they are synthetic constructs with GFP-encoding sequence inserted. An unusual group of recombinant enterovirus-G-like sequences, where the enterovirus structural protein-coding region has been switched for a porcine torovirus papain-like cysteine protease gene and other foreign sequences ^42,43^ were also discarded. ON457567 was removed because ∼230 codons in the 2B-2C region are shifted out of frame, explaining the high divergence of this sequence from other enterovirus sequences.

Next, we aimed to annotate the authentic polyprotein ORF (ppORF) in each sequence. In some sequences (particularly rhinoviruses) the longest AUG-to-stop-codon ORF begins 5′ of the authentic polyprotein initiation AUG codon (ppAUG). To annotate the correct ppAUG, we aligned sequences to the 30 RefSeq polyproteins. First, to check that the RefSeq annotations were themselves correct, we extracted the polyprotein amino acid sequences from the 30 RefSeqs and aligned them with MUSCLE version 3.8.31 ^44^. All 30 amino acid sequences aligned perfectly at the N terminus (i.e. no N-terminal alignment gaps), indicating that none had misannotated upstream initiation sites. Next, we identified the longest stop-codon-to-stop-codon ORF in each non-RefSeq sequence. Sequences that lack the ppORF stop codon or that have no ORF ≥ 3000 nt in length were removed. The corresponding amino acid sequences were aligned pairwise with each of the 30 RefSeq polyprotein sequences, again using MUSCLE. For each non-RefSeq sequence, the highest-identity RefSeq was determined. If the polyprotein initial methionine of this RefSeq aligned to a methionine in the non-RefSeq sequence, then that methionine was annotated as the N-terminus of the non-RefSeq polyprotein. The small number of remaining sequences were manually inspected via multiple sequence alignment with the 30 RefSeq, and all were found to be incomplete sequences that did not extend to the 5′ end of the ppORF; these sequences were therefore discarded. Sequences that were substantially truncated at the polyprotein C-terminus – namely sequences that did not align with the 30 RefSeqs at least up to a perfectly conserved “WH” ∼40 amino acids from the end of the polyprotein – were also discarded. We annotated the SL-VI region in each sequence by searching the 210 nt region immediately upstream of the ppAUG for the motif UU<u>AUG</u>GU[C/G]ACA, which is a (mostly) conserved sequence around the SL-VI AUG (underlined), or slight variations thereof. Sequences with >10 ambiguous nucleotide codes (for example ‘N’s) in the region from 10 nt upstream of the SL-VI AUG to the ppORF stop, sequences containing KEYWORDS tags UNVERIFIED, STANDARD_DRAFT, VIRUS_LOW_COVERAGE or VIRUS_AMBIGUITY in the GenBank files, and sequences with <160 nt of sequence upstream of the ppAUG, were discarded (for reference, the SL-VI AUG is 155, 152 and 157 nt upstream of the ppAUG in the enterovirus RefSeqs NC_001612, NC_001472 and NC_002058, respectively). At this stage, 9347 enterovirus sequences remained.

Next, we identified all AUG triplets within the 20 nt 5′ and 20 nt 3′ of the SL-VI AUG and calculated the coordinates of the corresponding ORFs, besides the SL-VI AUG ORF. There were 6137, 3165 and 45 sequences with 0, 1 or 2 AUGs (in addition to the SL-VI AUG) respectively; the SL-VI AUG itself was present in all except four of these sequences with the exceptions being AAG, AGG, AGG and AGG. In Lulla *et al.* (2019) ^5^, sequences were defined as having the enterovirus uORF if the ORF beginning with the SL-VI AUG and including the first in-frame stop codon (a) overlapped the ppORF by at least 1 nt, (b) was not in-frame with the ppORF, and (c) contained at least 150 nt upstream of the ppAUG. For each sequence, for the SL-VI AUG and any additional AUG triplets in the 20 nt 5′ and 20 nt 3′ flanking regions, we determined whether the associated ORF met the 2019 uORF criteria. Since there might be situations where a deletion in the uORF region has both reduced the distance between an upstream AUG and the ppAUG below the 150 nt threshold while also changing the reading frame so that UP expression depends on a non-SL-VI AUG, and also to allow for truncated versions of UP, we also investigated less stringent criteria for uORF definition, namely (a) encoded peptide ≥ 40 amino acids, (b) encoded peptide not contiguous with the polyprotein peptide, and (c) distance between the upstream AUG and the ppAUG ≥ 120 nt.

We used the single-linkage BLAST-based clustering algorithm BLASTCLUST ^45^ with parameters -p T -L 0.90 -b F -S 80 (i.e. ≥90% coverage of the shorter sequence, ≥80% amino acid identity threshold) to cluster the 9347 ppORF amino acid sequences, which resulted in 41 clusters. In each cluster, we chose a representative sequence for phylogenetic tree generation and tree annotation (Figure S3), as follows. If a cluster contained one or more NCBI RefSeqs, we used the RefSeq with the numerically lowest accession number. Otherwise, we looked for the ppORF amino acid sequence with the most identical copies (with ties broken arbitrarily) and, of the available corresponding accession numbers, we chose the one coming first alphanumerically. If there were no duplicated ppORF amino acid sequences, we chose the centroid sequence (minimum summed pairwise identity distances from ppORF amino acid sequence *i* to all other ppORF amino acid sequences *j* within the cluster). We aligned the 41 representative ppORF amino acid sequences with MUSCLE version 3.8.31 ^44^ and extracted the RdRp region (N-terminally trimmed to start at INAPSKTK-in NC_002058; this removes a region around the 3CPro-RdRp junction with many alignment gaps). A phylogenetic tree was estimated for the RdRp region using the Bayesian Markov chain Monte Carlo method implemented in MrBayes version 3.2.3, sampling across the default set of fixed amino acid rate matrices, with 5,000,000 generations, discarding the first 25% as burn-in (other parameters were left at defaults). The tree was visualized with FigTree version 1.4.2 (http://tree.bio.ed.ac.uk/software/figtree/). Transmembrane helix (TMH) domains and their confidence were predicted with Phobius (https://phobius.sbc.su.se/cgi-bin/predict.pl) ^10^.

### Cells and viruses

HEK293T cells (human embryonic kidney cell line, ATCC, CRL-3216), HeLa cells (ATCC, CCL-2) and HeLa Ohio cells (derivative of HeLa cell line, ECACC, 84121901) were maintained at 37°C in Dulbecco’s Modified Eagle Medium (DMEM, Lonza) supplemented with 10% fetal bovine serum (FBS), 1 mM L-glutamine, 20 mM HEPES (pH 7.3) and Penicillin/Streptomycin (10,000 U/ml). HIEC6 cells (human intestinal epithelial cell line, ATCC, CRL-3266) were maintained in Opti-MEM (Gibco) containing 20 mM HEPES, 1 mM L-Glutamine, Penicillin/Streptomycin (10,000 U/ml), 10 ng/ml hEGF (Sigma-Aldrich), and 5% FBS. All cells tested negative for mycoplasma. Human coxsackievirus A13 (CVA-13) strain Flores (EVAg, Ref-SKU: 014V-03623) was used for ribosome profiling and the development of a reverse genetics construct. Virus stocks were plaque purified and amplified using HeLa Ohio cells, clarified by centrifugation, purified through a 0.22 µm filter, and titrated on HeLa Ohio cells. Virus stocks were verified by sequencing virus-specific reverse transcribed PCR fragments, and used for ribosome profiling experiments.

### Plasmids

To create the reverse genetics clone for CVA-13, 5′ and 3′ terminal sequences (EVAg reference sequence AF465511.1) were used to amplify full-length genome using Phusion^TM^ High-Fidelity DNA polymerase (ThermoFisher Scientific). The amplified genome was cloned into vector pBR322 under the T7 promoter. The resulting clone was sequenced and deposited in GenBank (accession number PQ067515). The uuORF-KO mutation (AUG to GUG) was introduced using site-directed mutagenesis and confirmed by sequencing. The resulting plasmids were linearized with *Eag*I prior to T7 RNA transcription.

To evaluate the efficiency of IRES-mediated translation, a previously established pSGDLuc reporter cassette ^5^ was used to subclone terminal 5′ UTR sequences from CVA-13 (751 nt), CVA-1 (JX174177, 717 nt), and EV-A90 (JX390656, 751 nt). The cap-dependent *Renilla* luciferase gene was followed by an enterovirus 5′ UTR sequence and then, in one of the three frames (0, +1, +2), a 2A (StopGo) co-translational separation sequence followed by the firefly luciferase gene. The resulting plasmids were linearized with *Bam*HI prior to T7 RNA transcription. All plasmids were sequenced using the Plasmidsaurus full-plasmid sequencing service.

### RNA transcription

RNA transcription was performed using the mMESSAGE mMACHINE T7 transcription kit (Invitrogen). 10 µl transcription reactions were incubated at 37°C for 1 h, and reactions were terminated by treatment with DNase I for 15 min at 37°C. RNAs were further purified using the RNA Clean and Concentrator kit (Zymo Research) and quantified using Nanodrop.

### Recovery of CVA-13 virus from T7 transcripts

For virus recovery, 10^7^ HeLa Ohio cells were trypsinized, washed with PBS and electroporated with 20 µg T7 RNA in 800 µl PBS pulsed twice at 800 V and 25 µF using a Bio-Rad Gene Pulser Xcell^TM^ electroporation system. The cell suspension was supplemented with 10% FBS-containing media and incubated at 37°C. After 3 h of incubation and full cell attachment, the media was replaced with serum-free media, and cells were incubated until appearance of CPE. Virus stocks were amplified on HeLa Ohio cells using MOI 0.01, cleared by centrifugation, purified through a 0.22 µm filter, titrated on HeLa Ohio cells, and used for subsequent infections. All mutant viruses were also passaged at least 3 times at low MOI (0.01). The final virus stocks were used for RNA isolation, RT-PCR, and sequencing to confirm the presence of the introduced mutation.

### Reporter assay for relative IRES activity

HEK293T, HeLa and HIEC6 cells were transfected in triplicate using the method previously described ^5^. Briefly, per reaction, 100 ng of purified T7 transcribed RNA combined with 1 µl Lipofectamine 2000 (Invitrogen) in 10 µl Opti-Mem (Glibco) supplemented with RNaseOUT (Invitrogen; diluted 1:1,000 in Opti-Mem) were added to a suspension of cells at a density of 5×10^4^ cells (HeLa) or 1×10^5^ (HEK293T, HIEC6) per well on a 96-well plate. Once transfected, cells were supplemented with 5% FBS and incubated at 37°C for 8 h. Cells were lysed in 100 µl Passive Lysis Buffer (Promega) and freeze-thawed. Luciferase activity was measured using the Dual Luciferase Stop & Glo Reporter Assay System (Promega) as per manufacturer’s instructions. IRES-mediated translation was calculated as the ratio of IRES-dependent translation (firefly) to cap-dependent translation (*Renilla*). Translation in each of the three frames (0, +1, and +2) was then normalized to Frame 0 (corresponding to ppORF translation). Three independent experiments were performed.

### Virus infections and immunoblotting

HeLa Ohio cells at a density of 3.5×10^5^ per 35 mm plate were infected with CVA-13 at an MOI of 10 and incubated at 37°C for 0 to 18 h. Lysates were analyzed by SDS-PAGE, using standard 12% SDS-PAGE to resolve virus structural proteins and 4-20% Tris-Glycine gel (NuSep) to resolve UP as previously described ^5^. Structural proteins were detected using a pan-enterovirus monoclonal antibody (MA5-18206, Thermo Fisher) at 1:1,000 dilution, and cellular β-tubulin was detected using anti-tubulin antibody (ab15568, Abcam) at a 1:500 dilution. A custom rabbit polyclonal antibody raised against pre-TMH UP peptide CYHKANWIGHPVKVR (GenScript) was used at 1:200 dilution to detect CVA-13 UP. Immunoblots were imaged on a LI-COR ODYSSEY CLx imager and analyzed using Image Studio version 5.2.

### Culturing and infection of iPSC-derived neurons and human intestinal organoids

Preparation of iPSC-derived neurons was performed and validated as previously described ^16^. Infection was performed at the indicated MOI in cortical neuron media ^16^. After 1 h, the virus inoculum was removed and replaced with 50% fresh / 50% conditioned media. At the indicated times post-infection, media samples were taken and analyzed using a plaque assay in HeLa Ohio cells. For competition experiments, the media was replaced every other day to stimulate new virus release. Media samples were also used for RNA extraction, RT-PCR, and Sanger sequencing.

Human intestinal organoids were obtained from the duodenum of patients and grown as previously described ^5^. Differentiated monolayers cultured in 48-well plates were infected at the indicated MOI, and media samples were taken and analyzed by plaque assay in HeLa Ohio cells. For competition experiments, the media was replaced at 24 and 48 hpi to stimulate new virus release. Media samples were used for RNA extraction, RT-PCR, and Sanger sequencing. The differentiation of organoids was confirmed using RT-qPCR for the indicated cellular transcripts (Fig S7).

### Ribosome profiling and computational analysis of Ribo-Seq data

HeLa Ohio cells were grown on 150-mm dishes to a confluency of 80% and infected with CVA-13 at an MOI of 10 to ensure synchronous infection. For lactimidomycin (LTM) treated cells, 30 min before the specified time point, cells were treated with 50 mM LTM for 30 min. At 5 and 7 hpi, LTM treated and non-treated cells were flash frozen in an ethanol/dry ice bath and lyzed in the presence of 0.36 mM cycloheximide (CHX). The lysates were then processed according to previously described Ribo-Seq protocols ^5,13^. An Illumina NextSeq Platform was used to sequence the prepared amplicon libraries.

The bioinformatic Ribo-Seq analysis was performed as described previously ^5,13,15^. Briefly, adapter sequences were trimmed using the FASTX-Toolkit version 0.0.13 (http://hannonlab.cshl.edu/fastx_toolkit) and trimmed reads shorter than 25 nt were discarded. Reads were mapped to host (*Homo sapiens*) and virus RNA (CVA-13) using bowtie1 ^46^ version 0.12.947, with parameters -v 2 --best (i.e. maximum two mismatches, report best match). Mapping was performed in the following order: host rRNA, virus RNA, host RefSeq mRNA, host non-coding RNA, host genome. After calculating the read-length distributions (Fig. S5C), only 27–29 nt reads were taken forward for calculating phasing (Fig. S5D), and histograms of RPFs mapped to host mRNAs (Fig. S5A, S5B) and the viral genome (Fig. S5E, S5F). To normalize for library size in Figs S5E and S5F, reads per million mapped reads (RPM) values were calculated using the sum of positive-sense virus and host RefSeq mRNA 27–29 nt reads as the denominator. In Figs 4, S5E and S5F, a +12 nt offset was applied to the RPF 5′ end positions to give the approximate ribosomal P-site positions. To calculate the phasing (Fig. S5D) and length (Fig. S5C) distributions of host and virus RPFs, only RPFs whose 5′ end (+12 nt offset) mapped between the 13th nucleotide from the beginning and the 18th nucleotide from the end of coding sequences were counted; reads mapping to the dual coding region where the UP ORF overlaps the ppORF were also excluded. Histograms of host RPF 5′ end positions relative to initiation and termination sites (Figs S5A and S5B) were derived from RPFs mapping to RefSeq mRNAs with annotated coding regions ≥450 nt in length and with annotated 5′ and 3′ UTRs ≥60 nt in length.

For the analysis of host uAUG:mAUG ribosome occupancy ratios, we first selected a subset of host RefSeq mRNAs that have an upstream AUG codon (uAUG) within the annotated 5′ UTR. Initiation read counts for most host mRNAs were too low to obtain meaningful results for individual transcript species. Thus, in an attempt to obtain sufficient host mRNAs to average out sources of variation (shot noise, ligation bias, etc), we summed over uAUGs at a range of spacings from mAUGs (even though this might affect possible artifacts relating to stacking of preinitiation scanning ribosomes behind an initiating ribosome arrested at the mAUG ^14^). We avoided uAUGs very close to the 5′ end of annotated transcripts (at least 12 nt of sequence upstream of a uAUG are required for RPFs to map, but also very short 5′ UTRs are known to promote leaky scanning ^47^). While the CVA-13 uAUG is 157 nt 5′ of the mAUG, we allowed host uAUGs to be closer to the mAUG in order to increase the number of suitable host mRNAs. Furthermore, we reasoned that using uAUGs positioned closer to mAUGs may be more robust to concerns about alternative transcript isoforms when compared to more distally spaced uAUGs. Thus we selected host mRNAs with exactly one uAUG in the entire annotated 5′ UTR, and where that uAUG was within 200 nt upstream of the mAUG, and not in the first 30 nt of the annotated mRNA. This left 4684 out of an initial 35768 RefSeq mRNAs and ∼8% of all host mRNA-mapping RPFs. For both host and virus uAUG and mAUG ribosome occupancy levels, we again used only 27–29 nt reads, and counted only reads whose 5′ end (+12 nt offset) mapped to the A of the relevant AUG codon. For the histograms of host RPF 5′ end positions relative to uAUG and mAUG sites (Fig. S4), in contrast to Figs S5A and S5B, we removed the restriction that mRNAs should have annotated coding regions ≥450 nt in length and annotated 5′ and 3′ UTRs ≥60 nt in length.

## Acknowledgments

This work was funded by MRC project grant (MR/T000376/1) to V.L., M.Z. and A.E.F. V.L. is supported by a Sir Henry Dale Fellowship (220620/Z/20/Z) from the Wellcome Trust and the Royal Society and an Isaac Newton Trust/Wellcome Trust ISSF/University of Cambridge Joint Research Grant. A.E.F. is supported by a Wellcome Trust Senior Research Fellowship (220814/Z/20/Z). R.L.O. is supported by an MRC DTP Studentship. D.A.N. is supported by a Department of Pathology Gwynaeth Pretty PhD studentship. J.E.D. is supported by a Wellcome Trust Senior Research Fellowship (219447/Z/19/Z). The authors thank members of the Lulla lab for helpful discussions. We thank Cambridge Genomic Services for high-throughput sequencing.

The CVA-13 isolate (Flores) was provided by the European Virus Archive goes Global (EVAg) project funded by the European Union’s Horizon 2020 research and innovation programme under grant agreement No 871029.

For the purpose of Open Access, the authors have applied a CC BY public copyright license to any Author Accepted Manuscript (AAM) version arising from this submission.

