## Supplementary material for "Flexibility and modulation of translation initiation in enterovirus genomes"

| Number of sequences in the group | 20 nt 5' of SL-VI AUG ← | SL-VI AUG | → 20 nt 3' of SL-VI AUG | Representative sequence of the BLASTCLUST cluster to which these sequences belong |
|---|---|---|---|---|
| 1382 | ..........AUG....... | | .................... | NC_002058 Enterovirus C |
| 898 | ..........AUG....... | | .................... | NC_001430 Enterovirus D |
| 395 | ..........AUG....... | | .................... | NC_001612 Enterovirus A |
| 95 | .......AUG.......... | | .................... | NC_001612 Enterovirus A |
| 44 | .......AUG.......... | | .................... | MW969515 Rhinovirus C20 |
| 38 | .........AUG........ | | .................... | ON311169 Rhinovirus C15 |
| 35 | .......AUG.......... | | .................... | NC_001472 Enterovirus B |
| 34 | ..............AUG... | | .................... | NC_002058 Enterovirus C |
| 30 | ..............AUG... | | .................... | NC_038311 Rhinovirus A1 |
| 28 | ..........AUG....... | | .................... | NC_001472 Enterovirus B |
| 20 | .......AUGAUG....... | | .................... | NC_002058 Enterovirus C |
| 18 | .................... | | ...........AUG...... | ON881139 Rhinovirus C53 |
| 15 | .................... | | ..........AUG....... | NC_038311 Rhinovirus A1 |
| 12 | ..........AUG....... | | .................... | MW969532 Rhinovirus A101 |
| 11 | .................... | | ...........AUG...... | OL961531 Rhinovirus C13 |
| 9 | ..............AUG... | | .................... | NC_038878 Human rhinovirus NAT001 |
| 8 | ...........AUG...... | | ..........AUG....... | NC_001617 Rhinovirus A |
| 8 | ..............AUG... | | .................... | FJ445113 Rhinovirus A8 |
| 7 | .........AUG........ | | .................... | MW969515 Rhinovirus C20 |
| 7 | .........AUG........ | | .................... | NC_038311 Rhinovirus A1 |
| 7 | .......AUG.......... | | .................... | NC_002058 Enterovirus C |
| 7 | .........AUG........ | | .................... | NC_001617 Rhinovirus A |
| 6 | ..........AUG....... | | .................... | NC_038311 Rhinovirus A1 |
| 6 | ........AUG......... | | .................... | NC_002058 Enterovirus C |
| 5 | ............AUG..... | | .................... | NC_001490 Rhinovirus B14 |
| 4 | .................... | | .............AUG.... | NC_001859 Enterovirus E |
| 4 | ..............AUG... | | .................... | MW969532 Rhinovirus A101 |
| 4 | .................... | | ..........AUG....... | NC_001617 Rhinovirus A |
| 4 | ..............AUG... | | .................... | ON311169 Rhinovirus C15 |
| 4 | ..........AUG....... | | .................... | NC_038878 Human rhinovirus NAT001 |
| 3 | ...........AUG...... | | ...........AUG...... | ON311158 Rhinovirus C51 |
| 3 | .......AUGAUG....... | | .................... | NC_001612 Enterovirus A |
| 3 | .............AUG.... | | .................... | NC_001859 Enterovirus E |
| 3 | .................... | | ..........AUG....... | MW969532 Rhinovirus A101 |
| 3 | .................... | | ......AUG........... | NC_001472 Enterovirus B |
| 3 | ........AUG......... | | .................... | MZ835576 Rhinovirus C2 |
| 3 | .......AUG.......... | | .................... | LC316783 Enterovirus G |
| 2 | ...........AUGAUG... | | .................... | MW969532 Rhinovirus A101 |
| 2 | ..........AUG....... | | ..........AUG....... | NC_038311 Rhinovirus A1 |
| 2 | ..............AUG... | | ..............AUG... | NC_038311 Rhinovirus A1 |
| 2 | ......AUG.AUG....... | | .................... | NC_002058 Enterovirus C |
| 2 | ..........AUG.AUG... | | .................... | ON311169 Rhinovirus C15 |
| 2 | AUG................. | | .................... | NC_001859 Enterovirus E |
| 2 | .................... | | ......AUG........... | NC_038311 Rhinovirus A1 |
| 2 | .................... | | ............AUG..... | NC_038311 Rhinovirus A1 |
| 2 | .......AUG.......... | | .................... | HQ647168 Enterovirus A71 |
| 2 | .................... | | ...........AUG...... | NC_001472 Enterovirus B |
| 2 | .........AUG........ | | .................... | MW587060 Rhinovirus C |
| 2 | ...........AUG...... | | .................... | ON311158 Rhinovirus C51 |
| 2 | .......AUG.......... | | .................... | NC_004441 Porcine enterovirus 9 |
| 1 | ..............AUG... | | ..........AUG....... | NC_038311 Rhinovirus A1 |
| 1 | .........AUG........ | | .................... | NC_001859 Enterovirus E |
| 1 | .........AUG........ | | .................... | NC_001490 Rhinovirus B14 |
| 1 | .............AUG.... | | .................... | NC_001490 Rhinovirus B14 |
| 1 | .........AUG........ | | .................... | MW969532 Rhinovirus A101 |
| 1 | .............AUG.... | | .................... | KF688606 Rhinovirus C50 |
| 1 | ..........AUG....... | | .................... | AF326750 Enterovirus A125 |
| 1 | ..........AUG....... | | .................... | NC_029905 Enterovirus SEV-gx |
| 1 | ......AUG........... | | .................... | NC_001472 Enterovirus B |
| 1 | .................... | | ..............AUG... | NC_001472 Enterovirus B |
| 1 | ..........AUG....... | | .................... | MN865119 Enterovirus J |
| 1 | ..........AUG....... | | .................... | NC_010415 Enterovirus J |
| 1 | .......AUG.......... | | .................... | MZ835576 Rhinovirus C2 |
| 1 | .............AUG.... | | .................... | OK539488 Rhinovirus B27 |

**Supplementary Figure 1. Analysis of AUG triplets in regions flanking the SL-VI AUG in 9347 enterovirus sequences.** Columns show (1) the number of sequences with the given pattern of AUG triplets in the given BLASTCLUST cluster; (2) the positions of AUG triplets in the 20 nt 5′ of the SL-VI AUG; (3) the positions of AUG triplets in the 20 nt 3′ of the SL-VI AUG; and (4) the representative sequence of the relevant BLASTCLUST cluster (as in Figure S3). Note that different sequences in the same BLASTCLUST cluster may harbour different non-SL-VI AUG configurations, and so the representative sequence does not necessarily contain the displayed AUG configuration. Sequences containing any ambiguous nucleotide codes (e.g. "N", "R", etc) in the 20 nt 5′ or 20 nt 3′ of the SL-VI AUG, or sequences with incomplete coverage of this region, were removed. Sequences without any non-SL-VI AUG triplets in this region are also not shown. Note that the smallest distance between the SL-VI AUG and the ppAUG is 21 nt (in some rhinoviruses); thus none of the displayed AUG triplets corresponds to the ppAUG.

| | | | |
|---|---|---|---|
| #1 | MH933859 | Y | MAAYGDNLRLLSYSYWIGHPVTNRHIIYLFVGFVPLNSLSYQTLLYILKSNYLKQLWELKYLHRNQVHMRMQI |
| #1 | MH118028 | Y | MAAYGDNLRLLSYSYWIGHPVNSRDIIYLFVGFTKFNITSYTTLLYIILLNNKKRKWELKCLLRRPGLTRIKMLLLMVLP |
| #1 | MH118030 | Y | MAAYGDNLRLLSYSYWIGHPVNSRDIIYLFVGFTKLNITSYTTLLYIILLNNRKRKWELKCLLKRPDLTRTRMLLLTVLP |
| #1 | MH118031 | Y | MAAYGDNLRLLSYSYWIGHPVTSRDIIYLFVGFVKLDTTTLKSLLFITQLNIRKRKWELKSQLRRLDLTRIKT |
| #1 | AY697461 | Y | MAAYGDNLRLLSYSYWIGHPVTNRDLIYLFVGFTKLNITTFKTLLLIIQLNSEKRKWELKSQLRKLDRMRIKTSLQVDPL |
| #1 | MG253032 | Y | MAAYGDNLRLLSYSYWIGHPVTNRDIIYLFVGFIRLNTITFKTLLYIIQLNSRKRKWELKSQPKRLDLTRTKT |
| #1 | MG253033 | Y | MAAYGDNLRLLSYSYWIGHPVTNRDIIYLFVGFIRLNTITFKTLLYIIQLNSRKRKWELKSQPKRLDLMRTRT |
| #1 | MG253035 | Y | MAAYGDNLRLLSYSYWIGHPVTNRDIIYLFVGFIRLNTITFKTLLYIIQLNSRKGKWELKSQPKRLDLTRTRT |
| #1 | AY697459 | Y | MAAYGDNLRLLSYSYWIGHPVNSRDIIYLFVGFTTLNITNYRTLLYILLLNNRKRKWELKCPLKRPDRMRIRMLLPMALQ |
| #1 | KT277550 | Y | MAAYGDNLRLLSYSYWIGHPVNSRDIIYLFVGFTTLNITNHRTLLYILLLNNRKRKWELKCPLKRPGHMRIRTLLPMVLQ |
| #1 | AY697458 | Y | MAAYGDNLRLLSYSYWIGHPVNSRDIIYLFVGFIKLNITSYTTLLYIVLLNNRKRKWELKCLLRKPGLTRTKMLLLMVPP |
| #1 | AY773285 | Y | MAAYGDNLRLLSYSYWIGHPVTNRDLAYLFVGFTKLDIVTFKTLLYIIQLNSRKRKWELKFQPKRLDLMRTRT |
| #1 | JX390655 | Y | MAAYGDNLRLLSYSYWIGHPVNSRDIIYLFVGFTKLNITSYTTLLYIIQLNSEKRKWELKSQPKRLDLMRIRT |
| #1 | KU355877 | Y | MAAYGDNLRLLSYSFWIGHSVTNRDLVYLFVGFTKLNTITLKTLLLIIQLNSRKRKWELRYQPRKLDLMRTRISLRVDQL |
| #1 | JX390656 | Y | MAAYGDNLRLLSYSYWIGHPVNSRDIIYLFIGFTKLNITSYTTLLYIIQLNSRKRKWELKSQPKRLDLMRTRT |
| #1 | AB192877 | Y | MAAYGDNLRLLSYSYWIGHPVTNRDIIYLFVGFVKLDTTAFNSLLFITQLNNRKRKWELKSQPRKLDLTRTRT |
| #1 | AY697460 | Y | MAAYGDNLRLLSYSYWIGHPVTNRDIIYSFVGFVKLDSTTFKSLLFIIQLNNRKRKWELKSQPRKLDLTRTRT |
| #1 | JX390654 | Y | MAAYGDNLRLLSYSYWIGHPVNSRDIIYLFVGFTKFNITSYTTLLYIIQLNSRKRKWELKSQPKRLDLMRTRT |
| #1 | MG253034 | Y | MAAYGDNLRLLSYSYWIGHPVTNRDIIYLFVGFVRLNTITFKTLLYIIQLNSRKRKWELKSQPKRLDLTRTRT |
| #1 | ON809571 | Y | MAAYGDNLRLLSYSYWIGHPVTNKHIIYLFVGFVPLNTLSYQTLLYILGLNYSKQQWELRCPHRSQDRMKMRIWPLADLL |
| #1 | JF905564 | Y | MAAYGDNLRLLSYSYWIGHPVNSRDIIYLFVGFTKLNITSYTTLLYIILLNSKKRKWELKCLLKRPGLTRIRMLLLMVPP |
| #1 | MH118029 | Y | MAAYGDNLRLLSYSYWIGHPVNSRDIIYLFVGFTKLNITSYTTLLYIILLNNRKRKWELKCLLKRPDLTKTRMLLLTVLP |
| #2 | MN914206 | Y | MAAYGDNHRLLSYSDWIGHPVRKRDIIFLFVAFTPLNSFTPNLIKTVLLIRSVYHNGCASFITKGRGTRKHQCGYRWFNSKLHHY |
| #2 | AB828290 | N | MAAYGDNLRLLSYSYWIGHPVIYRDLRTNPPFYNTHLLINLIIHNGGSGVHSKVWIARKPECCCWWFYHQLHHYKLLQG |
| #2 | KX932039 | N | MAAYGDNLRLLSYSCWIGHPVIYRDLRTNPPFYNTHLLINLTIQNGSSSVHSEIWITREPESCCWRFHHQLYHHKLLQRQC |
| #2 | KC785523 | N | MLMVTIVLISCCHYSRSVCNYKNKIISYYHPSRSVHNYKNEIFCHNGCSGICSEQWNTRKQKHSH |
| #2 | MF990294 | N | MAAYGDNLRLLSYSIWIGHPVISKATHLSYCFTVFIYLLHPSNHNGSSSVHSEIWITREPECCSWWIHY |
| #2 | MT432142 | N | MAAYGDNQRLLPQSELDWPSGECCVRYTTVCWNHCVSFTSHLTN |
| #2 | OK570211 | Y | MAAYGDNLRLLSYSNWIGHPVRVRDIVYLFVAFAPLNTYTPSLIRIILLIRSAYHNGCTSFISKGRSP |
| #2 | OK570194 | Y | MAAYGDNLGLLSYSNWIGHPVRVRDIIYLFVAFTPLNTYTPNLIKTVLLIRSLYHNGCTSFISEGGST |
| #2 | OM963010 | N | MAAYGDNLRLLSYSCWIGHPVNCRDLNIYSFLCITYSLNISVQP |
| #2 | LS451300 | Y | MAAYGDNLGLLSYSNWIGHPVRVRDIIYLFVAFTPLDTNTPILIRAVILIRSIHHNGCTSFISKGGST |
| #2 | OK570210 | Y | MAAYGDNLRLLSYSNWIGHPVRIRDIIYLFVAYTPLNSYTPDLIKVVLLIRSIHHNGCASFISEGGST |
| #2 | LS451301 | Y | MAAYGDNLRLLSYSDWIGHPVRIRDIIYLFVAFTPLNSYTPSLIKAVLLIRSAYHNGCTGFISKGGST |
| #2 | ON383157 | Y | MAAYGDNLRLLSYSCWIGHPVKVKDIVYLFVAFTPLNNRTLSLIKIVLLLRAIYHNGCTGFIPKGGGT |
| #2 | KC344834 | N | MAAYGDNQRLLSYSFWIGHPVIFETITFTCLTTQLSTLLHSHSHNGSSSVHTKVWFS |
| #2 | KC785524 | N | MLMVTIVLISCCHCSRSVCNYENKTISCYHPPRSVHNHKNKIFCHNGCSGICSEQWNPRKQKHSHWWFNHQLHYY |
| #2 | OP410421 | N | MAAYGDNQRLLSYSELDWPSGECCVRYTTVCWNHCVSFTSHLTN |
| #2 | MG571859 | N | MAAYGDNLRLLSYSYWIGHPVIFESDICLAISLHLLTYNTANSQWELKCPLRNLDHMRIKTWLLVVPLLITPP |
| #2 | MZ092702 | N | MLMVTIVLISCCHCSHSVCNHENKTISCYHPSRSVYNHKNEFFCHNGCSGICSEQWNPRKQKHSHWWLNHQLHYY |
| #2 | MK250423 | N | MAAYGDNLRLLSYSYWTGHPVILKVILALLFHCICLFTILQTHNGSSSVHSEIRIT |
| #2 | AF499635 | N | MAAYGDNLRLLSYSFWIGHPVIFEINIPSLLLHSTHLFITLL |
| #2 | JX174176 | N | MAAYGDNLRLLSYSVWIGHPVTLKAIYLTHCLTEFTYLLHLYNNNGSSSVYSEVWIT |
| #2 | JX174177 | N | MAAYGDNLRLLSYSVWIGHPVTSKAIYLTHCLTEFTYLLHLYNNNGSSSVYSEVWIT |
| #2 | PP461545 | Y | MAAYGDNLRLLSYSYWIGHPVTNRDLIYLFVGFTKLNITNFKTLLFIIQLNNRKRKWELKYHHRKWVLMRTLM |
| #2 | KJ170436 | Y | MWLLMVTITDCYHKANWIGHPVKVRFIIYLFAGFAPLSVFTLSTISTVISIRQLYHNGCSGFITESGRT |
| #2 | KJ170510 | Y | MLMVTITDCYHKANWIGHPVKVRFIIYLFAGFAPLSVFTLSTISTVISIRQLYHNGCSGFITESGRT |
| #2 | AF499641 | N | MAAYGDNLGLLSYSFWIGHPVTSNVQVVYSLNFLNTFSVVEL |
| #2 | KR815824 | N | MLMVTIVLISCCHCSHSVCNYKNKTISCYHPSRSVYNHKNEFFCYNGCSGICSE |
| #2 | JX982253 | N | MLMVTIVLISCCQYSRSVCNYKNKIISYYNPSRSVHNYENEIFCHNGCSGICSEQWNTRKQKHSHWWFNH |
| #2 | JX982254 | N | MLMVTIVLISCCHYSRSVCNYKNKIISYYNPSRSVHNYENEIFCHNGCSGICSEQWNTRKQKHSHWWFNH |
| #2 | JX982255 | N | MLMVTIVLISCCHYSRSVCNYKNKIISYYNPSRSVHNYENEIFCHNGCSGICSEQWNTRKQKHSHWWFNH |
| #2 | JX982256 | N | MLMVTIVLISCCHYSRSVCNYKNKIISYYHPSRSVHNYKNEIFCHNGCPGICSEQWNTRKQKHSHWWFNH |
| #2 | JX982257 | N | MLMVTIVLISCCHYSRSVCNYKNKIISYHHPSRSVHNYKNEIFCHNGCSGICSEQWNTRKQKHSHWWFNH |
| #2 | JX982258 | N | MLMVTIVLISCCHYSRSVCNYKNKIISYYNPSRSVHNYENEIFCHNGCSGICSEQWNTRKQKHSHWWFNH |
| #2 | JX982259 | N | MLMVTIVLISCCHYSRSVCNYKNKIISYYHPSRSVHNYKNEIFCHNGCSGICSEQWNTRKQKHSHWWFNH |
| #2 | AB686524 | N | MLMVTIVLISCCHCSRSVCNYKNKTISCYHPPRSVHNHKNEIFCHNGRSSICSEQWNPRKQKHSHWWFNHQLHYY |
| #2 | MZ092704 | N | MLMVTIVLISCCHCSHSVCNHENKTISCYHPSRSAYNHKNEFFCHNGCSGICSEQWNPRKQKHSHWWLNHQLHYY |
| #2 | MH484166 | Y | MWLLMVTIIGCYHKANWIGHPVRVKHIIYLFVGFTPLSVFTPDVIRVVLLIRSFYHNGSSSIIPEGRST |
| #2 | OP137321 | N | MWLLMVTIIDCYHKANWIGHPVKYKHIIYLLVGFISLTNLPLA |
| #2 | JX275107 | Y | MAAYGNNHRLLSKSELDWPSSVNQINYSLVCWIRSRNVLLFNLLKLFEDRILVSQWELKYHPKK |
| #2 | EF015017 | Y | MWLLMVTIIDCYHKANWIGHPVKYKHLIYLLVGFTPLTQFTPSIISTVLLVRHYHRYGCTGLISESGRS |
| #2 | EF015015 | Y | MWLLMVTIIDCYHKASWIGHPVKYKHIIYLFVGFTPLTQFTPSIIIIVLLIRHQYQYGRTGLISESGCS |
| #2 | EF015012 | Y | MAAYGDNLRLLSYSNWIGHPVRVRDIIYLFVAFTPLSTNTVTLIRIVLLIRLTHHYGCTSFISEGGCT |
| #2 | EF555644 | Y | MAAYGDNLGLLSYSDWIGHPVRIKDIVYLFVAFTPLNKNTLSLIKVVLLIRSAYHNGCTSLISEGRGTRKHQRGHRWFNRKLHYHQLLQRFG |
| #2 | EF015030 | Y | MAAYGAITDCYHKANWIGHPVKYKHIIYIFVGFTLLTHSSPKLIYTVLVIRNYHFSYGSSSIHSKDWGPREPECGCKRVHNQLYNHQLL |
| #3 | KX981987 | N | MERLLPYSYWIGHPVTNRAIIYLFVRFIPLSLKEVKTLQFIVKLNTAKWELKYQRKRLGHMRPG |
| #25 | AF414373 | N | MAAYGDNLTLLPYSCWVGHPDFVIKPFPRALTFNTFVNFFKSLLLVLKY |
| #40 | AF326750 | Y | MAVYGDNLEYSDCHHKLLGLANRSIIYYFVGFVPLTPIIYKILLRFLLANLRYQRWVHKFLHNNLELMKTLMLQQEDQPYTTPPSTTTKTLMQHQQTSKIFHRTLQSLHNQWWMP |

**Supplementary Figure 2. Non-SL-VI AUG ORFs with length at least 40 codons.** Amino acid sequences of all ORFs that start with a non-SL-VI AUG within the 20 nt 5′ and 20 nt 3′ of the SL-VI AUG and that fulfill the following criteria: (a) encoded peptide ≥ 40 amino acids, (b) encoded peptide not contiguous with the polyprotein peptide, and (c) distance between the non-SL-VI AUG and the ppAUG ≥ 120 nt. The sequence WIGHP that is conserved in some enterovirus UP proteins is highlighted in blue. Transmembrane helix (TMH) predictions are highlighted in red (20–50% confidence), yellow (50–80% confidence) or green (>80% confidence); underlined sequences represent predicted N-terminal signal peptides (Phobius predictions). Columns show (1) BLASTCLUST cluster; (2) NCBI accession; (3) 'Y' if the non-SL-VI AUG ORF meets the more stringent Lulla *et al.* (2019) [1] uORF criteria (i.e. if the ORF beginning at the AUG and including the first in-frame stop codon overlaps the ppORF by at least 1 nt; is not in-frame with the ppORF; and contains at least 150 nt upstream of the ppAUG), otherwise 'N'; and (4) the amino acid sequence of the ORF. For the purpose of cross-referencing with Figure S3, the 'representative sequences' for the relevant clusters are: #1 NC_001612 Enterovirus A, #2 NC_002058 Enterovirus C, #3 NC_001472 Enterovirus B, #25 NC_010415 Enterovirus J, and #40 AF326750 Enterovirus A125. Note that some of these ORFs also have an in-frame SL-VI AUG (e.g. EF015017 – here, the SL-VI AUG corresponds to the methionine at position 5).

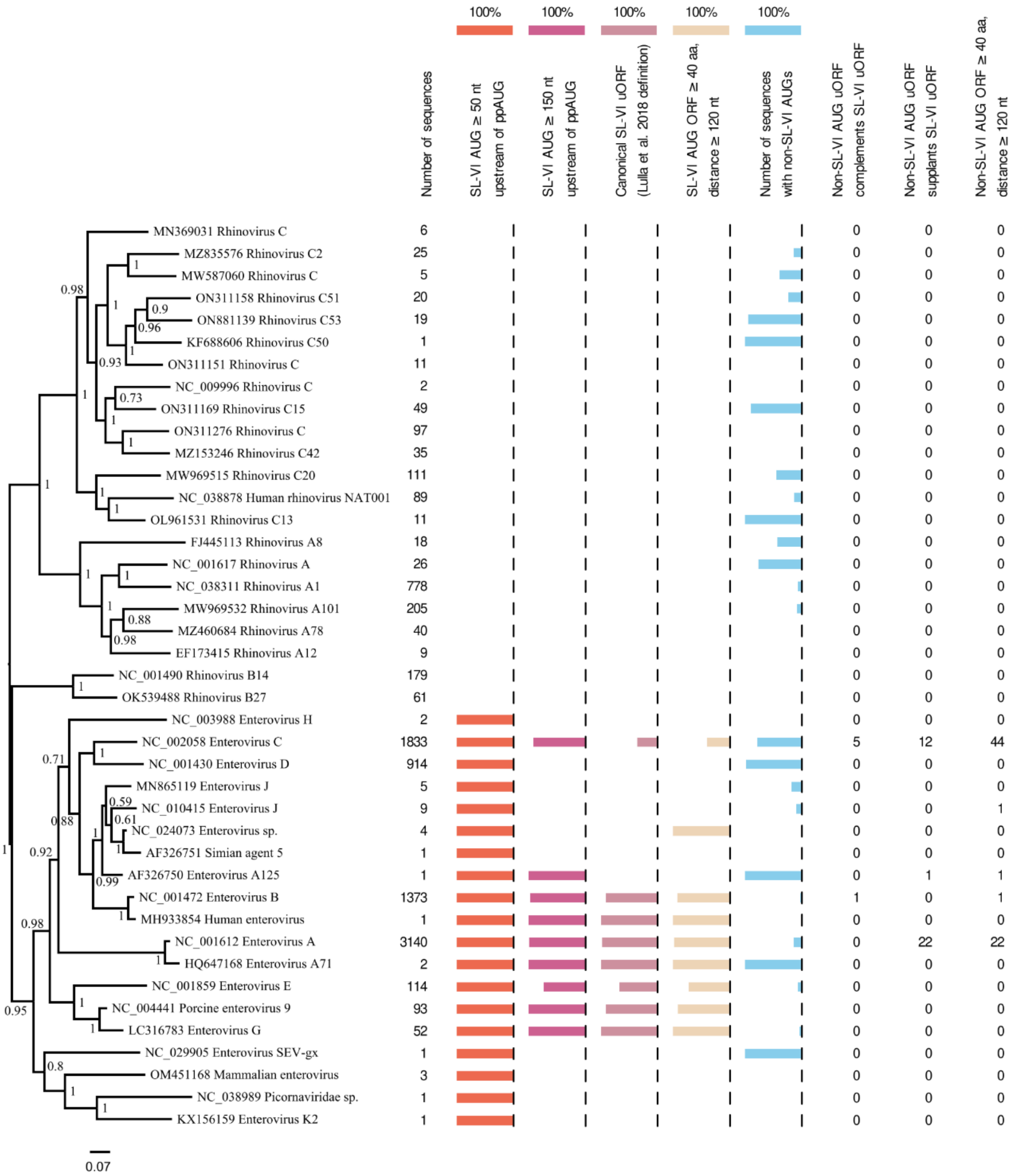


**Supplementary Figure 3. uORF statistics for different clusters of enterovirus sequences.** The ppORF amino acid sequences of 9347 enterovirus sequences were clustered with BLASTCLUST using an 80% identity threshold, a representative sequence was selected from each of the 41 clusters, the ppORF amino acid sequences were aligned with MUSCLE, and a phylogenetic tree (left) was estimated with MrBayes (see Methods). Upstream AUG and upstream ORF statistics were calculated for each cluster. "distance ≥ 120 nt" refers to the distance between the upstream AUG and the ppAUG. "Non-SL-VI AUG uORF complements SL-VI uORF" indicates that the non-SL-VI AUG uORF and the SL-VI AUG uORF share the same termination codon [note, in the one case in the NC_001472 cluster, the non-SL-VI AUG is 3 codons 3′ of the SL-VI AUG and the resulting uORF fails the Lulla *et al.* (2019) [1] uORF criterion that the uORF initiation codon should be at least 150 nt upstream of the ppAUG]. "Non-SL-VI AUG uORF supplants SL-VI uORF" indicates that the non-SL-VI AUG uORF fulfills the Lulla *et al.* (2019) [1] uORF criteria, but the ORF beginning with the SL-VI AUG does not.

A

| | repeat | time point | host uAUGs | host mAUGs | ratio (u/m) |
|---|---|---|---|---|---|
| NT | 1 | 5 h p.i. | 67 | 145 | 0.46 |
| NT | 2 | 5 h p.i. | 70 | 178 | 0.39 |
| NT | 1 | 7 h p.i. | 2 | 6 | 0.33 |
| NT | 2 | 7 h p.i. | 4 | 19 | 0.21 |
| LTM | 1 | 5 h p.i. | 693 | 2306 | 0.30 |
| LTM | 2 | 5 h p.i. | 1079 | 3371 | 0.32 |
| LTM | 1 | 7 h p.i. | 20 | 79 | 0.25 |
| LTM | 2 | 7 h p.i. | 15 | 58 | 0.26 |

B

| | repeat | time point | virus uAUG | virus mAUG | ratio (u/m) |
|---|---|---|---|---|---|
| NT | 1 | 5 h p.i. | 121 | 162 | 0.75 |
| NT | 2 | 5 h p.i. | 139 | 98 | 1.4 |
| NT | 1 | 7 h p.i. | 67 | 37 | 1.8 |
| NT | 2 | 7 h p.i. | 337 | 90 | 3.7 |
| LTM | 1 | 5 h p.i. | 9582 | 922 | 10.4 |
| LTM | 2 | 5 h p.i. | 17521 | 1566 | 11.2 |
| LTM | 1 | 7 h p.i. | 735 | 253 | 2.9 |
| LTM | 2 | 7 h p.i. | 427 | 140 | 3.1 |

C

| | repeat | time point | uAUG context | host uAUGs | host mAUGs | ratio (u/m) |
|---|---|---|---|---|---|---|
| LTM | 1 | 5 h p.i. | all | 693 | 2306 | 0.30 |
| LTM | 1 | 5 h p.i. | poor | 185 | 725 | 0.26 |
| LTM | 1 | 5 h p.i. | medium | 333 | 1120 | 0.30 |
| LTM | 1 | 5 h p.i. | strong | 175 | 461 | 0.38 |
| LTM | 2 | 5 h p.i. | all | 1079 | 3371 | 0.32 |
| LTM | 2 | 5 h p.i. | poor | 333 | 1108 | 0.30 |
| LTM | 2 | 5 h p.i. | medium | 521 | 1620 | 0.32 |
| LTM | 2 | 5 h p.i. | strong | 225 | 643 | 0.35 |

D

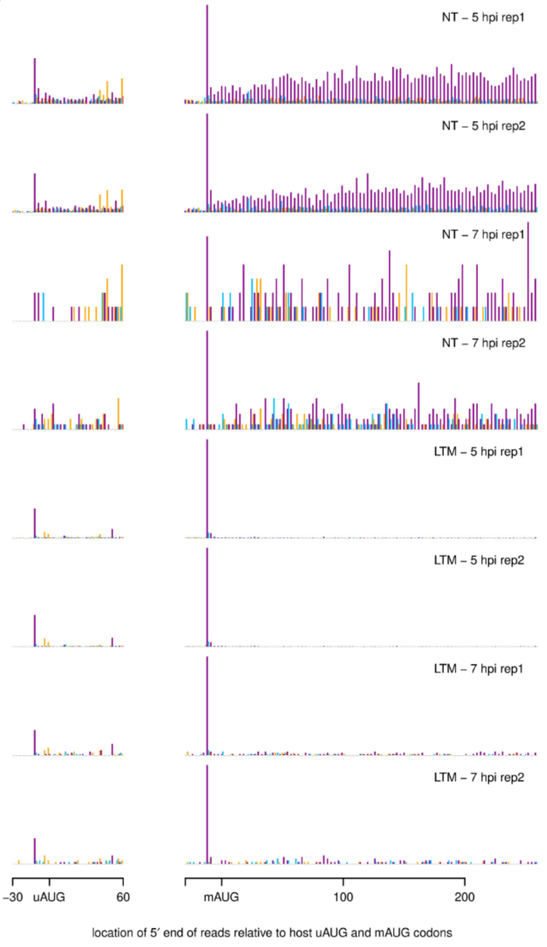


**Supplementary Figure 4. Relative RPF occupancy at upstream and main AUG codons.** (**A**) After first mapping RPFs to host RefSeq mRNAs, those mapping to annotated transcripts with exactly one upstream AUG (uAUG) and where the uAUG was within 200 nt upstream of the main AUG (mAUG) and not in the first 30 nt of the transcript, were selected. Next, RPFs whose 5′ end (+12 nt offset) mapped to the A of the uAUG or mAUG were quantified. Due to host-cell shut-off and the limited number of suitable host mRNAs used, the numbers of uAUG- and mAUG-mapping RPFs were often small, but useful numbers were obtained at least for the 5 h p.i. LTM libraries (marked in red). (**B**) As for panel A but for the CVA-13 virus uAUG and mAUG codons. (**C**) Data from the 5 h p.i. LTM libraries in panel A, subdivided by uAUG initiation context: all contexts, strong contexts (G at −3 and +4, or A at −3), medium contexts (G at −3 or G at +4), and poor contexts (other contexts). As expected, there is a modest increase in uAUG:mAUG occupancy ratios with increasing strength of the uAUG initiation context. (**D**) Histograms of the 5′-end mapping positions of RPFs (no +12 nt offset applied) relative to uAUG and mAUG codons, summed over the selected set of host mRNAs. RPFs whose 5′ ends map to the 1st, 2nd or 3rd positions of codons are shown in purple, blue or yellow, respectively. In all panels, only 27-29 nt reads were used.

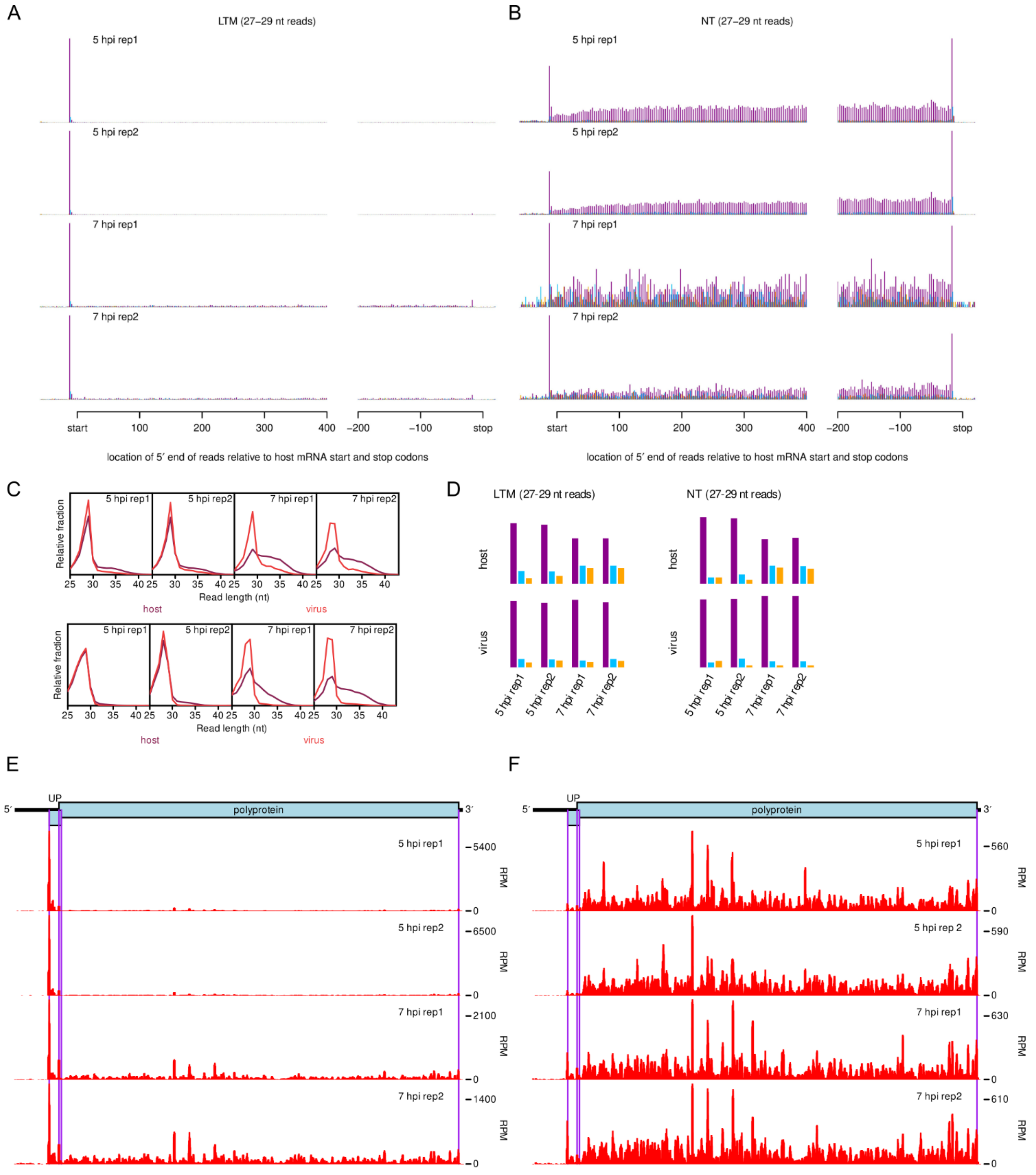


**Supplementary Figure 5. Assessment of ribosome profiling quality for lactimidomycin-treated (LTM) and non-treated (NT) CVA-13 libraries.** (**A-B**) Histograms of the 5′-end mapping positions of RPFs relative to annotated initiation and termination sites, summed over all host mRNAs. Reads whose 5′ ends map to the 1st, 2nd or 3rd positions of codons are shown in purple, blue or yellow, respectively. (**C**) Length distributions for Ribo-Seq reads mapping to virus (red) or host mRNA (purple) coding regions; upper panels – LTM, lower panels – NT. (**D**) Phasing of 5′ ends of Ribo-Seq reads that map to the virus or host mRNA coding regions. (**E-F**) Ribosome profiles for the CVA-13 genome at 5 and 7 hpi for cells treated with lactimidomycin (E) or untreated (F). Histograms of the 5′-end mapping positions of RPFs, with a +12 nt offset to map the approximate P-site position, in reads per million mapped reads (RPM), smoothed with a 15-nucleotide running mean filter. For panels A, B, D, E and F, only 27–29 nt reads were used.

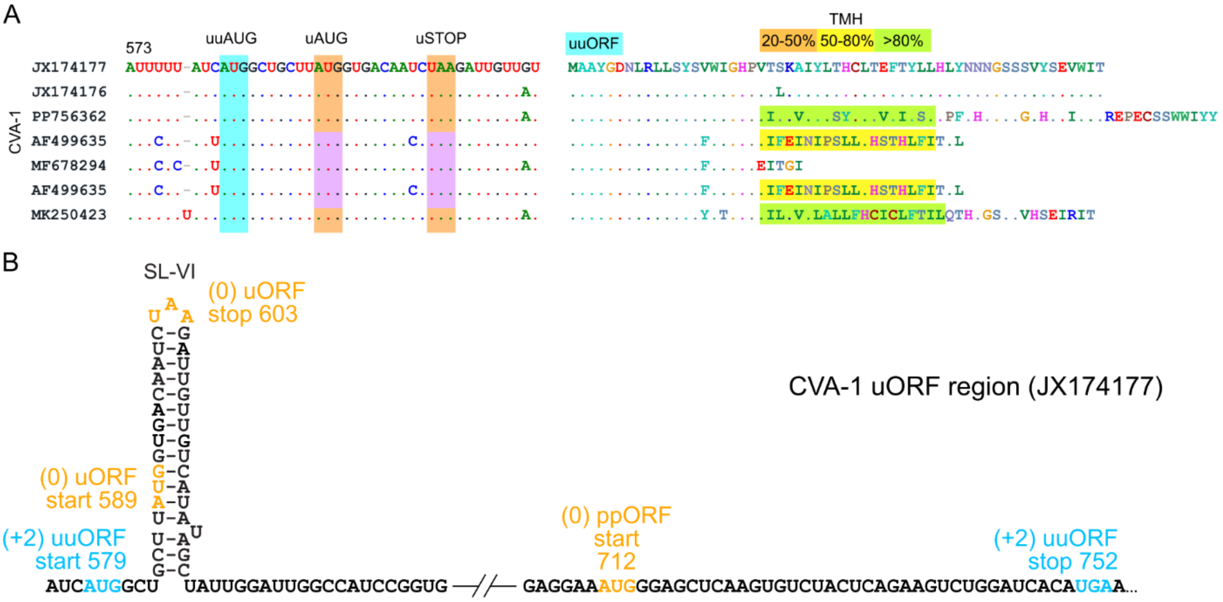


**Supplementary Figure 6. Enterovirus CVA-1 sequences that encode a UP-like protein from an alternative upstream AUG codon.** (**A**) Nucleotide (left) and uuORF-encoded protein (right) sequences in enterovirus CVA-1, where the uORF is truncated and the uuORF can potentially rescue UP expression. Transmembrane helix (TMH) predictions are highlighted in yellow (50–80% confidence) or green (>80% confidence). Depending on the frame used, ORFs are highlighted in blue (uuORF), and purple or orange (uORF). (**B**) Schematic representation of the CVA-1 IRES dVI region with the uuORF (blue), uORF and ppORF (orange) start and stop codons annotated.

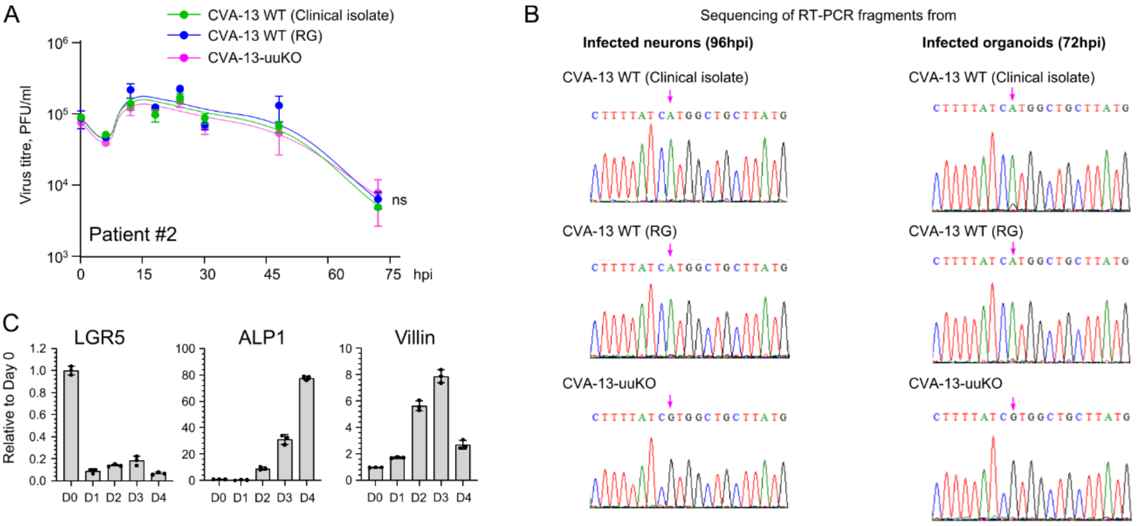


**Supplementary Figure 7. Validation of organoid and neuron infection experiments.** (**A**) The growth of CVA13 viruses in organoid line derived from patient 2. (**B**) Representative sequencing chromatograms of RT-PCR products from viruses derived from the final time points in Fig. 7A and 7C. (**C**) Differentiation of human intestinal organoid cultures. Results of qRT-PCR showing fold change in transcript levels in differentiated duodenum organoids (days 1-4) relative to transcript levels in undifferentiated organoids (day 0). The GAPDH transcript was used for normalization. LGR5, a stem cell marker, leucine-rich repeat-containing G-protein coupled receptor 5; ALP, a mature enterocyte marker, alkaline phosphatase; Villin, an epithelial cell marker. Plotted data represent means ± s.d.; n = 3.

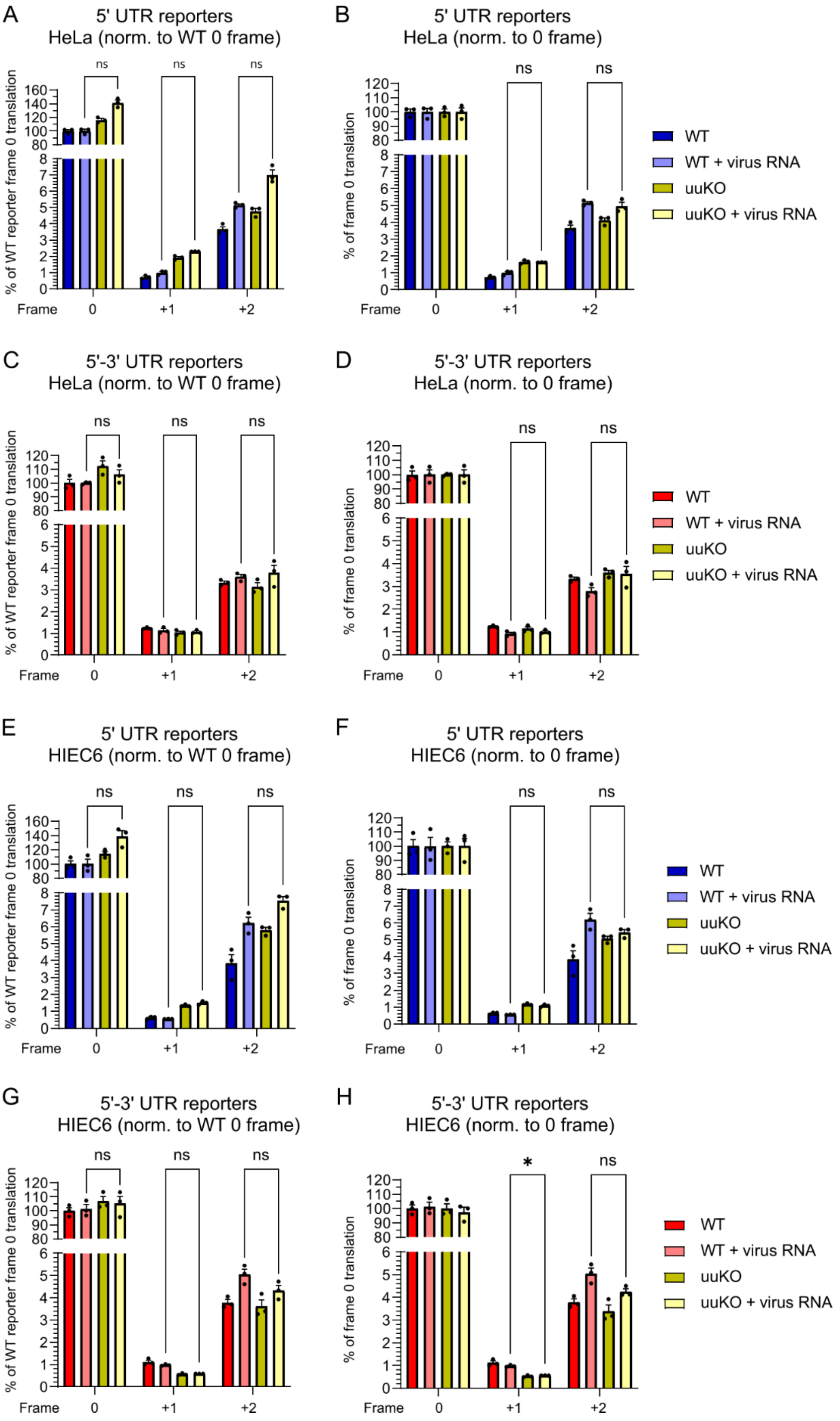


**Supplementary Figure 8. Full set of expression assays performed for 5′UTR and 5′-3′UTR reporters.** Analysis of IRES activities for CVA-13 5′-3′ UTR reporters in the three frames in HeLa and intestinal HIEC6 cells, with and without virus RNA, at 8 h.p.t. Statistical analysis was conducted using two-tailed *t*-tests; * $p$ value ≤ 0.05; ns, non significant.

| #seqs | +0 frame relative to SL-VI AUG | +1 frame relative to SL-VI AUG | +2 frame relative to SL-VI AUG |
|---|---|---|---|
| 5123 | ... ... ... ... ... ... AUG ... ... ... ... ... ... ... ... ... ... | . ... ... ... ... ... ... ... ... ... ... ... ... ... ... ... ... ... .. | .. ... ... ... ... ... ... ... ... ... ... ... ... ... ... ... ... ... . |
| 1276 | ... ... ... ... ... ... AUG ... ... ... ... ... ... ... ... ... ... | . ... ... ... ... ... ... ... ... ... ... ... ... ... ... ... ... ... .. | .. ... ... ... AUG nnn nnn nnn nnn nnn nnn nnn nnn nnn nnn nnn STP ... . |
| 933 | ... ... ... ... ... ... AUG nnn nnn nnn STP ... ... ... ... ... ... | . ... ... ... ... ... ... ... ... ... ... ... ... ... ... ... ... ... .. | .. ... ... ... AUG ... ... ... ... ... ... ... ... ... ... ... ... ... . |
| 397 | ... ... ... ... ... ... AUG nnn nnn nnn nnn nnn STP ... ... ... ... | . ... ... ... ... ... ... ... ... ... ... ... ... ... ... ... ... ... .. | .. ... ... ... ... ... ... ... ... ... ... ... ... ... ... ... ... ... . |
| 363 | ... ... ... ... ... ... AUG ... ... ... ... ... ... ... ... ... ... | . ... ... ... ... ... ... ... ... ... ... ... ... ... ... ... ... ... .. | .. ... ... ... AUG nnn nnn nnn nnn nnn nnn STP ... ... ... ... ... ... . |
| 187 | ... ... ... ... ... ... AUG nnn nnn nnn nnn nnn nnn nnn nnn nnn STP | . ... ... ... ... ... ... ... ... ... ... ... ... ... ... ... ... ... .. | .. ... ... ... ... ... ... ... ... ... ... ... ... ... ... ... ... ... . |
| 99 | ... ... ... ... ... ... AUG nnn nnn nnn nnn nnn nnn STP ... ... ... | . ... ... ... ... ... ... ... ... ... ... ... ... ... ... ... ... ... .. | .. ... ... ... ... ... ... ... ... ... ... ... ... ... ... ... ... ... . |
| 86 | ... ... ... ... ... ... AUG ... ... ... ... ... ... ... ... AUG ... | . ... ... ... ... ... ... ... ... ... ... ... ... ... ... ... ... ... .. | .. ... ... ... ... ... ... ... ... ... ... ... ... ... ... ... ... ... . |
| 79 | ... ... ... ... ... ... AUG nnn nnn nnn nnn nnn nnn nnn nnn STP ... | . ... ... ... ... ... ... ... ... ... ... ... ... ... ... ... ... ... .. | .. ... ... ... ... ... ... ... ... ... ... ... ... ... ... ... ... ... . |
| 77 | ... ... ... ... ... ... AUG ... ... ... ... ... ... ... ... ... ... | . ... ... ... ... ... ... ... ... ... ... ... ... ... ... ... ... ... .. | .. ... ... AUG nnn nnn nnn nnn nnn nnn nnn STP ... ... ... ... ... ... . |
| 68 | ... ... ... ... ... ... AUG ... ... ... ... ... ... ... ... ... ... | . ... ... ... ... ... ... ... ... ... ... ... ... ... ... ... ... ... .. | .. ... ... ... AUG nnn nnn nnn nnn nnn nnn nnn STP ... ... ... ... ... . |
| 58 | ... ... ... ... ... ... AUG ... ... ... ... ... ... ... ... ... ... | . ... ... ... ... ... ... ... ... ... ... ... ... ... ... ... ... ... .. | .. ... ... AUG nnn nnn nnn nnn nnn nnn nnn nnn nnn nnn nnn nnn nnn STP . |
| 58 | ... ... ... ... ... ... AUG nnn nnn nnn STP ... ... ... ... ... ... | . ... ... ... ... ... ... ... ... ... ... ... ... ... ... ... ... ... .. | .. ... ... ... ... ... ... ... ... ... ... ... ... ... ... ... ... ... . |
| 52 | ... ... ... ... AUG ... AUG ... ... ... ... ... ... ... ... ... ... | . ... ... ... ... ... ... ... ... ... ... ... ... ... ... ... ... ... .. | .. ... ... ... ... ... ... ... ... ... ... ... ... ... ... ... ... ... . |
| 47 | ... ... ... ... ... ... AUG ... ... ... ... ... ... ... ... ... ... | . ... ... ... AUG nnn nnn nnn nnn STP ... ... ... ... ... ... ... ... .. | .. ... ... ... ... ... ... ... ... ... ... ... ... ... ... ... ... ... . |
| 43 | ... ... ... ... ... ... AUG nnn nnn nnn nnn nnn STP ... ... ... ... | . ... ... ... ... ... ... ... ... ... ... ... ... ... ... ... ... ... .. | .. ... ... AUG ... ... ... ... ... ... ... ... ... ... ... AUG ... ... . |
| 43 | ... ... ... ... ... ... AUG nnn nnn nnn nnn STP ... ... ... ... ... | . ... ... ... ... ... ... ... ... ... ... ... ... ... ... ... ... ... .. | .. ... ... ... ... ... ... ... ... ... ... ... ... ... ... ... ... ... . |
| 41 | ... ... ... ... ... ... AUG ... ... ... ... ... ... ... ... ... ... | . ... ... ... ... ... ... ... ... ... ... ... ... ... ... ... ... ... .. | .. ... ... ... AUG ... ... ... ... ... ... ... ... ... ... ... ... ... . |
| 37 | ... ... ... ... ... ... AUG ... ... ... ... ... ... ... ... ... ... | . ... ... ... ... ... ... ... ... ... ... ... ... ... ... ... ... ... .. | .. ... ... ... AUG nnn nnn nnn nnn nnn nnn nnn nnn nnn nnn nnn nnn STP . |
| 21 | ... ... ... ... ... ... AUG nnn nnn nnn nnn nnn STP ... ... ... ... | . ... ... ... ... ... ... ... ... ... ... ... ... ... ... ... ... ... .. | .. ... ... ... ... ... ... ... ... ... ... ... ... ... ... AUG ... ... . |
| 20 | ... ... ... ... AUG nnn AUG nnn nnn nnn nnn nnn nnn nnn nnn STP ... | . ... ... ... ... ... ... ... ... ... ... ... ... ... ... ... ... ... .. | .. ... ... ... ... ... ... ... ... ... ... ... ... ... ... ... ... ... . |
| 19 | ... ... ... ... ... ... AUG ... ... ... ... ... ... ... ... ... ... | . ... ... ... ... ... ... ... ... ... ... ... ... ... ... ... ... ... .. | .. ... ... AUG AUG nnn nnn nnn nnn nnn nnn nnn nnn nnn nnn nnn STP ... . |
| 18 | ... ... ... ... ... ... AUG nnn nnn nnn nnn nnn nnn STP ... ... ... | . ... ... ... ... ... ... ... ... ... ... ... ... ... ... ... ... ... .. | .. ... ... ... ... ... ... ... ... ... ... ... AUG nnn nnn nnn nnn STP . |
| 15 | ... ... ... ... ... ... AUG ... ... ... ... ... ... ... ... ... ... | . ... ... ... ... ... ... ... ... ... ... ... AUG nnn nnn nnn STP ... .. | .. ... ... ... ... ... ... ... ... ... ... ... ... ... ... ... ... ... . |
| 14 | ... ... ... ... AUG nnn AUG nnn nnn nnn nnn STP ... ... ... ... ... | . ... ... ... ... ... ... ... ... ... ... ... ... ... ... ... ... ... .. | .. ... ... ... ... ... ... ... ... ... ... ... ... ... ... ... ... ... . |
| 10 | ... ... ... ... ... ... AUG ... ... ... ... ... ... ... AUG ... ... | . ... ... ... ... ... ... ... ... ... ... ... ... ... ... ... ... ... .. | .. ... ... ... ... ... ... ... ... ... ... ... ... ... ... ... ... ... . |
| 9 | ... ... ... ... ... ... AUG ... ... ... ... ... ... ... ... ... ... | . ... ... ... ... ... ... ... ... ... ... ... ... ... ... ... ... ... .. | .. ... ... AUG nnn nnn nnn nnn nnn nnn nnn nnn nnn nnn nnn nnn STP ... . |
| 9 | ... ... AUG ... ... ... AUG ... ... ... ... ... ... ... ... ... ... | . ... ... ... ... ... ... ... ... ... ... ... ... ... ... ... ... ... .. | .. ... ... ... ... ... ... ... ... ... ... ... ... ... ... ... ... ... . |
| 9 | ... ... ... ... ... ... AUG ... ... ... ... ... ... ... ... ... ... | . ... ... ... ... ... ... ... ... ... ... ... ... ... ... ... AUG ... .. | .. ... ... ... ... ... ... ... ... ... ... ... ... ... ... ... ... ... . |
| 8 | ... ... ... AUG nnn nnn AUG nnn nnn nnn nnn nnn nnn STP ... ... ... | . ... ... ... ... ... ... ... ... ... ... ... AUG ... ... ... ... ... .. | .. ... ... ... ... ... ... ... ... ... ... ... ... ... ... ... ... ... . |
| 7 | ... ... ... ... ... ... AUG nnn nnn nnn nnn nnn nnn STP ... ... ... | . ... ... ... AUG nnn nnn nnn nnn STP ... ... ... ... ... ... ... ... .. | .. ... ... ... ... ... ... ... ... ... ... ... ... ... ... ... ... ... . |
| 7 | ... ... ... ... ... ... AUG nnn nnn nnn nnn STP ... ... ... ... ... | . ... ... ... ... ... ... ... ... ... ... ... ... ... ... ... ... ... .. | .. ... ... ... ... ... ... ... ... ... ... ... AUG ... ... ... ... ... . |
| 6 | ... ... ... ... ... ... AUG nnn nnn nnn nnn nnn STP ... ... ... ... | . ... ... ... AUG nnn nnn nnn nnn nnn nnn nnn STP ... ... ... ... ... .. | .. ... ... ... ... ... ... ... ... ... ... ... ... ... ... AUG ... ... . |
| 4 | ... ... ... ... ... ... AUG nnn nnn nnn nnn STP ... ... ... ... ... | . ... ... ... ... ... ... ... ... ... ... ... ... ... ... ... ... ... .. | .. ... ... ... ... ... ... ... ... ... ... ... AUG nnn nnn nnn STP ... . |
| 4 | ... ... ... ... ... ... AUG nnn nnn nnn nnn nnn nnn STP ... ... ... | . ... ... ... ... ... ... ... ... ... ... ... AUG ... ... ... ... ... .. | .. ... ... ... ... ... ... ... ... ... ... ... ... ... ... ... ... ... . |
| 4 | ... ... ... ... ... ... AUG ... ... ... ... ... ... ... ... ... ... | . ... ... ... ... ... ... ... ... ... ... ... ... AUG ... AUG ... ... .. | .. ... ... ... ... ... ... ... ... ... ... ... ... ... ... ... ... ... . |
| 4 | ... ... ... ... ... ... ... ... ... ... ... ... ... ... ... ... ... | . ... ... ... ... ... ... ... ... ... ... ... ... ... ... ... ... ... .. | .. ... ... ... ... ... ... ... ... ... ... ... ... ... ... ... ... ... . |
| 3 | ... ... ... ... ... ... AUG nnn nnn nnn nnn nnn nnn nnn nnn STP ... | . ... ... ... ... ... ... ... ... ... ... ... ... ... ... ... ... ... .. | .. ... ... ... AUG ... ... ... ... ... ... ... ... ... ... ... ... ... . |
| 3 | ... ... ... ... ... ... AUG ... ... ... ... ... ... ... ... ... ... | . ... ... ... ... ... ... ... ... ... ... ... ... ... ... ... ... ... .. | .. ... ... AUG AUG nnn nnn nnn nnn nnn nnn STP ... ... ... ... ... ... . |
| 3 | ... ... ... ... ... ... AUG ... ... ... ... ... ... ... ... ... ... | . ... ... ... ... ... ... ... ... ... ... ... ... ... ... ... ... ... .. | .. ... ... ... ... AUG nnn nnn nnn nnn nnn nnn STP ... ... ... ... ... . |
| 3 | ... ... ... ... ... ... AUG nnn nnn nnn nnn nnn STP ... ... ... ... | . ... ... ... AUG ... ... ... ... ... ... ... ... ... ... ... ... ... .. | .. ... ... ... ... ... ... ... ... ... ... ... ... ... ... AUG ... ... . |
| 3 | ... ... ... ... ... ... AUG nnn nnn nnn nnn STP ... ... ... ... ... | . ... ... ... ... AUG ... ... ... ... ... ... ... ... ... ... ... ... .. | .. ... ... ... ... ... ... ... ... ... ... ... ... ... ... ... ... ... . |
| 3 | ... ... ... ... ... ... AUG ... ... AUG ... ... ... ... ... ... ... | . ... ... ... ... ... ... ... ... ... ... ... ... ... ... ... ... ... .. | .. ... ... ... ... ... ... ... ... ... ... ... ... ... ... ... ... ... . |
| 3 | ... ... ... ... ... ... AUG ... ... ... ... ... ... ... ... ... ... | . ... ... ... ... ... ... ... ... ... ... ... AUG ... ... ... ... ... .. | .. ... ... ... ... ... ... ... ... ... ... ... ... ... ... ... ... ... . |
| 3 | ... ... ... ... ... ... AUG ... ... ... ... ... ... ... ... ... ... | . ... ... ... ... ... ... ... ... ... ... ... ... ... ... ... ... AUG .. | .. ... ... ... ... ... ... ... ... ... ... ... ... ... ... ... ... ... . |
| 2 | ... ... ... ... ... ... AUG ... ... ... ... ... ... ... ... ... ... | . ... ... AUG nnn nnn nnn nnn nnn STP ... ... ... ... ... ... ... ... .. | .. ... ... ... AUG nnn nnn nnn nnn nnn nnn nnn nnn nnn nnn nnn STP ... . |
| 2 | ... ... ... ... ... ... AUG nnn nnn nnn nnn nnn nnn nnn nnn STP ... | . ... ... ... ... ... ... ... ... ... ... ... ... ... ... ... ... ... .. | .. ... ... ... AUG nnn nnn nnn nnn nnn nnn nnn STP ... ... ... ... ... . |
| 2 | ... ... ... ... ... ... AUG nnn nnn nnn STP ... ... ... ... ... ... | . ... ... ... ... ... ... ... ... ... ... ... ... ... ... ... ... ... .. | .. ... ... ... AUG nnn nnn nnn nnn nnn nnn nnn nnn nnn nnn nnn STP ... . |
| 2 | ... ... ... AUG nnn nnn AUG nnn nnn nnn nnn STP ... ... ... ... ... | . ... ... ... ... ... ... ... ... ... ... ... ... ... ... ... ... ... .. | .. ... ... ... ... ... ... ... ... ... ... ... AUG nnn nnn nnn nnn STP . |
| 2 | ... ... ... ... ... ... AUG ... ... ... ... ... ... ... ... ... ... | . ... ... ... ... ... ... ... ... ... ... ... AUG nnn nnn nnn STP ... .. | .. ... ... ... AUG nnn nnn nnn nnn nnn nnn nnn STP ... ... ... ... ... . |
| 2 | ... ... ... ... AUG nnn AUG nnn nnn nnn nnn nnn nnn nnn nnn STP ... | . ... ... ... ... ... ... ... ... ... ... ... ... ... ... ... ... ... .. | .. ... ... ... ... ... ... ... ... ... ... ... ... AUG ... ... ... ... . |
| 2 | ... ... ... ... ... ... AUG nnn nnn nnn nnn nnn nnn nnn nnn nnn STP | . ... ... ... ... ... ... ... ... ... ... ... ... ... ... ... ... ... .. | .. ... ... ... ... AUG ... ... ... ... ... ... ... ... ... ... ... ... . |
| 2 | ... ... ... ... AUG ... AUG ... ... ... ... ... ... ... ... ... AUG | . ... ... ... ... ... ... ... ... ... ... ... ... ... ... ... ... ... .. | .. ... ... ... AUG nnn nnn nnn nnn nnn nnn nnn nnn STP ... ... ... ... . |
| 2 | ... ... ... ... ... ... AUG nnn nnn nnn nnn STP ... ... ... ... ... | . ... ... ... ... ... ... ... ... ... ... ... ... ... ... ... ... ... .. | .. ... ... ... ... ... ... ... ... ... ... ... AUG nnn nnn nnn nnn STP . |
| 2 | ... ... ... ... ... ... AUG nnn nnn nnn nnn nnn nnn AUG nnn STP ... | . ... ... ... ... ... ... ... ... ... ... ... ... ... ... ... ... ... .. | .. ... ... ... ... ... ... ... ... ... ... ... ... ... ... ... ... ... . |
| 2 | ... ... ... ... ... ... AUG ... ... ... ... ... ... ... ... ... ... | . AUG nnn nnn nnn nnn nnn nnn nnn STP ... ... ... ... ... ... ... ... .. | .. ... ... ... ... ... ... ... ... ... ... ... ... ... ... ... ... ... . |
| 2 | ... ... ... ... ... ... AUG nnn nnn AUG nnn nnn STP ... ... ... ... | . ... ... ... ... ... ... ... ... ... ... ... ... ... ... ... ... ... .. | .. ... ... ... ... ... ... ... ... ... ... ... ... ... ... ... ... ... . |
| 2 | ... ... ... AUG AUG ... AUG ... ... ... ... ... ... ... ... ... ... | . ... ... ... ... ... ... ... ... ... ... ... ... ... ... ... ... ... .. | .. ... ... ... ... ... ... ... ... ... ... ... ... ... ... ... ... ... . |
| 2 | ... ... ... AUG ... ... AUG ... ... ... ... ... ... ... ... ... ... | . ... ... ... ... ... ... ... ... ... ... ... ... ... ... ... ... ... .. | .. ... ... ... ... ... ... ... ... ... ... ... ... ... ... ... ... ... . |
| 2 | ... ... ... ... AUG ... AUG ... ... ... ... ... ... ... ... ... AUG | . ... ... ... ... ... ... ... ... ... ... ... ... ... ... ... ... ... .. | .. ... ... ... ... ... ... ... ... ... ... ... ... ... ... ... ... ... . |
| 2 | ... ... ... ... ... ... AUG ... ... ... ... ... ... ... ... ... ... | . ... ... ... ... AUG ... ... ... ... ... ... ... ... ... ... ... ... .. | .. ... ... ... ... ... ... ... ... ... ... ... ... ... ... ... ... ... . |
| 1 | ... ... ... ... ... ... AUG nnn nnn nnn nnn nnn nnn nnn nnn STP ... | . ... ... ... ... ... ... ... ... ... ... ... ... ... ... ... ... ... .. | .. ... ... ... AUG nnn nnn nnn nnn nnn nnn STP ... ... ... ... ... ... . |
| 1 | ... ... ... AUG nnn nnn AUG nnn nnn nnn nnn nnn nnn STP ... ... ... | . ... ... ... ... ... ... ... ... ... ... ... ... ... ... ... ... ... .. | .. ... ... ... ... ... ... ... ... ... ... ... AUG nnn nnn nnn nnn STP . |
| 1 | ... ... ... ... ... ... AUG ... ... ... ... ... ... ... ... ... ... | . ... ... ... ... ... ... ... ... ... ... ... ... ... ... ... ... ... .. | .. ... ... AUG nnn nnn nnn nnn nnn nnn nnn nnn nnn nnn STP ... ... ... . |
| 1 | ... ... ... ... ... ... AUG nnn nnn nnn nnn nnn nnn nnn nnn nnn STP | . ... ... ... AUG ... ... ... ... ... ... ... ... ... ... ... ... ... .. | .. ... ... ... ... ... ... ... ... ... ... ... ... ... ... ... ... ... . |
| 1 | ... ... ... ... AUG nnn AUG nnn nnn nnn nnn nnn nnn nnn STP ... ... | . ... ... ... ... ... ... ... ... ... ... ... AUG ... ... ... ... ... .. | .. ... ... ... ... ... ... ... ... ... ... ... ... ... ... ... ... ... . |
| 1 | ... ... ... ... ... ... AUG ... ... ... ... ... ... ... ... ... ... | . ... ... ... ... ... ... ... ... ... ... ... ... ... ... ... ... ... .. | .. ... ... ... AUG nnn nnn nnn nnn nnn nnn nnn nnn STP ... ... ... ... . |
| 1 | ... ... ... ... AUG nnn AUG nnn nnn nnn nnn nnn nnn STP ... ... ... | . ... ... ... ... ... ... ... ... ... ... ... ... ... ... ... ... ... .. | .. ... ... ... ... ... ... ... ... ... ... ... ... ... ... ... ... ... . |
| 1 | ... ... ... ... ... ... AUG nnn nnn nnn nnn nnn nnn nnn STP ... ... | . ... ... ... ... ... ... ... ... ... ... ... ... ... ... ... ... ... .. | .. ... ... ... ... ... ... ... ... ... ... ... ... ... ... ... ... ... . |
| 1 | ... ... ... ... ... ... AUG nnn nnn nnn nnn AUG nnn nnn nnn STP ... | . ... ... ... ... ... ... ... ... ... ... ... ... ... ... ... ... ... .. | .. ... ... ... ... ... ... ... ... ... ... ... ... ... ... ... ... ... . |
| 1 | ... ... ... ... ... ... AUG ... ... ... ... ... ... ... ... ... ... | . ... ... ... ... ... ... ... ... ... ... ... ... ... ... ... ... ... .. | .. ... ... AUG AUG nnn nnn nnn nnn nnn nnn nnn STP ... ... ... ... ... . |
| 1 | ... ... ... ... ... ... AUG nnn nnn nnn nnn nnn nnn STP ... ... ... | . ... ... ... ... ... ... ... ... ... ... ... ... ... ... ... ... ... .. | .. ... ... ... ... ... ... ... ... ... ... ... ... ... ... ... AUG ... . |
| 1 | ... ... ... ... ... ... AUG ... ... ... ... ... ... ... ... ... ... | . ... ... AUG nnn nnn nnn nnn nnn STP ... ... ... ... ... ... ... ... .. | .. ... ... ... ... ... ... ... ... ... ... ... ... ... ... ... ... ... . |
| 1 | ... ... ... ... ... ... AUG nnn nnn nnn nnn STP ... ... ... ... ... | . ... ... ... ... ... ... ... ... ... ... ... ... ... ... ... ... ... .. | .. ... ... ... AUG ... ... ... ... ... ... ... ... ... ... ... ... ... . |
| 1 | ... ... ... ... ... ... AUG nnn nnn nnn STP ... ... AUG ... ... ... | . ... ... ... ... ... ... ... ... ... ... ... ... ... ... ... ... ... .. | .. ... ... AUG STP ... ... ... ... ... ... ... ... ... ... ... ... ... . |
| 1 | ... ... ... ... ... ... AUG nnn nnn nnn STP ... ... AUG ... ... ... | . ... ... ... ... ... ... ... ... ... ... ... ... ... ... ... ... ... .. | .. ... ... ... ... ... ... ... ... ... ... ... ... ... ... ... ... ... . |
| 1 | ... ... ... ... ... ... AUG nnn nnn nnn STP ... ... ... ... ... ... | . ... ... ... ... ... ... ... ... ... ... ... ... ... ... ... ... ... .. | .. ... ... ... ... AUG ... ... ... ... ... ... ... ... ... ... ... ... . |
| 1 | ... ... ... ... ... ... AUG ... ... ... ... ... ... ... ... ... ... | . ... ... ... ... ... ... ... ... ... ... ... ... ... ... ... ... ... .. | .. ... ... ... ... ... ... ... ... ... ... ... ... AUG nnn nnn nnn STP . |
| 1 | ... ... ... ... ... ... AUG ... ... ... ... ... ... ... ... ... ... | . ... ... ... ... ... ... ... ... ... ... ... ... ... ... ... ... ... .. | .. ... ... ... ... ... ... ... ... ... ... ... ... ... ... AUG nnn STP . |
| 1 | ... ... ... ... ... ... AUG ... ... ... ... AUG ... ... ... ... ... | . ... ... ... ... ... ... ... ... ... ... ... ... ... ... ... ... ... .. | .. ... ... ... ... ... ... ... ... ... ... ... ... ... ... ... ... ... . |
| 1 | ... ... ... ... ... ... AUG ... ... ... ... ... ... ... ... ... ... | . ... ... ... ... ... ... ... ... ... ... ... ... ... ... ... ... ... .. | .. ... ... ... ... ... ... ... ... ... ... ... ... ... ... AUG ... ... . |
| 1 | ... ... ... ... ... ... AUG ... ... ... ... ... ... ... ... ... ... | . ... ... ... ... ... ... ... ... ... ... ... ... ... ... ... ... ... .. | .. ... ... ... ... ... ... ... ... ... ... ... ... ... ... ... ... AUG . |
| 1 | ... ... ... ... ... ... AUG STP ... ... ... ... ... ... ... ... ... | . ... ... ... ... ... ... ... ... ... ... ... ... ... ... ... ... ... .. | .. ... ... ... ... ... ... ... ... ... ... ... ... ... ... ... ... ... . |

**Supplementary Figure 9. Mapping positions of short ORFs in SL-VI region.** A region from 20 nt 5′ of the SL-VI AUG to 30 nt 3′ of the SL-VI AUG was extracted. All AUG codons are shown in the three reading frames. Stop codons (STP) are only shown if they terminate an ORF initiated by one (or more) of the displayed AUGs. Other nucleotides are shown as "n" if they are part of one of these ORFs, otherwise as ".". Sequences with incomplete coverage of this region, or any ambiguous nucleotide codes ("R", "N", etc) in this region were removed, leaving 9333 sequences. Number of sequences (#seqs) list the number of sequences with the given configuration of displayed AUGs, stops, and short ORFs.

**Supplementary Table 1**. Metadata associated with *E. alphacoxsackie* species.

| **Enterovirus A isolate sequence** | **Type** | **Isolation source** | **Isolation country** | **Year** | **Associated disease** | **Reference PMID** |
|---|---|---|---|---|---|---|
| KT277550 | A89 | stool | China | 2011 | contact of an AFP patient | 26685900 |
| AY697459 | A89 | stool | Bangladesh | 2000 | isolated from an AFP patient | 15659764 |
| MH118030 | A76 | N/A | India | 2011 | N/A | N/A |
| MH118029 | A76 | N/A | India | 2010 | N/A | N/A |
| MH118028 | A76 | N/A | India | 2010 | N/A | N/A |
| AY697458 | A76 | stool | Bangladesh | 1999 | isolated from an AFP patient | 15659764 |
| JF905564 | A76 | stool | China | 2004 | isolated from an AFP patient | 21755310 |
| MH118031 | A90 | N/A | India | 2010 | N/A | N/A |
| MH933859 | A90 | stool | Cameroon | 2014 | N/A | N/A |
| AB192877 | A90 | N/A | Cambodia | 2005 | isolated from an AFP patient | N/A |
| AY773285 | A90 | N/A | Netherlands | 2004 | isolated from an AFP patient | N/A |
| AY697460 | A90 | stool | Bangladesh | 1999 | isolated from an AFP patient | 15659764 |
| JX390654 | A90 | stool | China | 2001 | isolated from an AFP patient | 23081679 |
| JX390655 | A90 | stool | China | 2001 | isolated from an AFP patient | 23081679 |
| JX390656 | A90 | stool | China | 2003 | isolated from an AFP patient | 23081679 |
| MG253035 | A90 | stool | China | 2011 | isolated from an AFP patient | 29980696 |
| MG253034 | A90 | stool | China | 2011 | isolated from an AFP patient | 29980696 |
| MG253033 | A90 | stool | China | 2011 | isolated from an AFP patient | 29980696 |
| MG253032 | A90 | stool | China | 2011 | isolated from an AFP patient | 29980696 |
| KU355877 | A121 | stool | India | 2013 | healthy child | 27902407 |
| AY697461 | A91 | stool | Bangladesh | 2000 | isolated from an AFP patient | 15659764 |
| ON809571 | A119 | raw wastewater | France | 2015 | N/A | 37212710 |
| AF326750 | A125 | baboon stool | | 1962 | N/A | 11773400 |

AFP – acute flaccid paralysis, N/A – no data available.

**Supplementary Table 2.** Metadata associated with *E. coxsackiepol* species.

| Enterovirus C isolate sequence | Type | Isolation source | Isolation country | Year | Associated disease | Reference PMID |
|---|---|---|---|---|---|---|
| EF015012 | EV-C99 | clinical isolate, CDC | Oklahoma, USA | 1985 | N/A | 19264596 |
| EF015030 | CVA-21 | stool | Bangladesh | 2000 | AFP | 19264596 |
| EF555644 | EV-C99 | clinical isolate, CDC | Georgia, USA | 1984 | N/A | 19264596 |
| JX275107 | PV2 | PV2 outbreak | Nigeria | 2008 | paralytic disease (poliomyelitis) | 23408630 |
| LS451300 | EV-C99 | stool | Madagascar | 2003 | polio surveillance | 30323802 |
| LS451301 | EV-C99 | stool | Madagascar | 2003 | polio surveillance | 30323802 |
| MN914206 | EV-C99 | stool | Malawi | 2003 | surveillance | 32629843 |
| OK570194 | EV-C99 | stool | Madagascar | 2011 | polio surveillance | 36348312 |
| OK570210 | EV-C99 | stool | Madagascar | 2011 | polio surveillance | 36348312 |
| OK570211 | CVA-20 | stool | Madagascar | 2011 | polio surveillance | 36348312 |
| ON383157 | EV-C99 | stool | Guatemala | 2020 | environmental water from human excretion | 35950869 |
| PP461545 | CVA-24 | stool | Nepal | 2023 | N/A | N/A |

AFP – acute flaccid paralysis, N/A – no data available.

**Supplementary Table 3. Host and virus read counts for Ribo-Seq samples.**

| treatment | repeat | time point | total reads | host mRNA(+) | vRNA(+) | host mRNA(+) | vRNA(+) |
|---|---|---|---|---|---|---|---|
| | | | | all reads ≥25 nt | | 27–29 nt reads | |
| NT | 1 | 5 h p.i. | 7,816,493 | 337,247 | 202,634 | 182,022 | 142,629 |
| NT | 2 | 5 h p.i. | 6,651,515 | 393,300 | 296,507 | 220,144 | 226,696 |
| NT | 1 | 7 h p.i. | 11,437,523 | 32,125 | 23,185 | 13,893 | 16,891 |
| NT | 2 | 7 h p.i. | 9,203,363 | 102,111 | 85,105 | 40,605 | 63,126 |
| LTM | 1 | 5 h p.i. | 8,427,129 | 190,159 | 72,899 | 114,910 | 53,860 |
| LTM | 2 | 5 h p.i. | 11,635,282 | 278,943 | 107,787 | 162,110 | 75,024 |
| LTM | 1 | 7 h p.i. | 9,384,830 | 46,725 | 22,484 | 16,822 | 13,186 |
| LTM | 2 | 7 h p.i. | 7,860,105 | 35,459 | 20,473 | 13,951 | 12,975 |
